# Spatiotemporal Orchestration of Mitosis by Cyclin-Dependent Kinase

**DOI:** 10.1101/2024.04.29.591629

**Authors:** Nitin Kapadia, Paul Nurse

**Author notes:** Correspondence to Nitin Kapadia.

## Abstract

Mitotic onset is a critical transition for eukaryotic cell proliferation. The prevailing view for its control is that cyclin-dependent kinase (CDK) is activated first in the cytoplasm, at the centrosome, which then initiates mitosis^1–3^. Bistability in CDK activation ensures the transition is irreversible but how this unfolds in a spatially compartmentalized cell is unknown^4–8^. Here using fission yeast, we show that CDK is actually activated in the nucleus first, not the cytoplasm, and that the bistable responses dramatically differ within the nucleus and cytoplasm. There is a stronger response in the nucleus permitting mitotic signal propagation from there to the cytoplasm. Abolishing cyclin-CDK localization to the yeast centrosome led to activation occurring only in the nucleus, spatially uncoupling the nucleus and cytoplasm mitotically, suggesting centrosomal cyclin-CDK acts as a “signal relayer”. We propose that the key mitotic regulatory system operates in the nucleus in proximity to DNA, allowing incomplete DNA replication and DNA damage to be effectively monitored to preserve genome integrity and for integration of ploidy within the CDK control network. This spatiotemporal regulatory framework establishes core principles for control of the onset of mitosis *in vivo*, which will help inform how CDK controls mitotic onset in other eukaryotes.

## Main

Entry to mitosis is a complex cellular transition. The cell undergoes numerous rapid changes across the cell, including modifications to the nuclear membrane, chromosome condensation, disassembly of cytoskeletal microtubules, and formation of the mitotic spindle^9^. These changes are mainly driven by cyclin-CDK complexes^10^ through phosphorylation of hundreds of different substrates^10–12^. However, a full understanding of the spatial and temporal dynamics of the CDK control network and how these bring about mitosis has been difficult to establish. This is in part due to there being multiple different cyclin-CDK complexes and regulators in most eukaryotes, which have potential overlapping and redundant functions, making interpretation of experiments complex^12–17^. In addition, *in vitro* systems designed to investigate the dynamics of CDK activation while taking account of cellular spatial compartmentation, have been limited and may not completely represent the dynamics present *in vivo*^5, 6, 16^. Finally, there is a lack of precise and simultaneous measurements of *in vivo* CDK activity and endogenous levels of CDK and its regulators^18, 19^.

The Cdk1 kinase in complex with its regulatory subunit, cyclin, is the main driver triggering cellular changes at mitotic entry^20^. Its activity gradually rises through the cell cycle until late G2, when it rises more rapidly to bring about the onset of mitosis, through a combination of cyclin accumulation and regulation of the phosphorylation of a Cdk1 tyrosine residue (Y15 in most eukaryotes) near its ATP binding site^4–6, 21, 22^. The inhibitor Wee1 kinase family phosphorylates Y15 to inactivate Cdk1, while the activator Cdc25 family dephosphorylates this site ^23–27^. Cdk1 also phosphorylates Wee1 and Cdc25, respectively inactivating and activating them^28–31^. Theoretical modelling and biochemical experiments of these positive (Cdk1 and Cdc25) and double-negative (Cdk1 and Wee1) feedback loops have shown they are able to abruptly increase Cdk1 activity at mitotic entry^4–6^. Furthermore, these feedback loops give rise to systems-level features of bistability and hysteresis, where for a given concentration of cyclin, CDK activity can either be high if the state of the system is mitotic or low if it is in interphase^4, 6, 8^. This ensures the mitotic transition is irreversible and that the interphase and mitotic states are clearly distinguished, thus preventing cells slipping back and forth between them^4–6, 32^.

This regulation is complicated by the fact that the eukaryotic cell contains multiple spatial compartments, the most prominent being the nucleus and cytoplasm. Such spatial complexity poses a challenge to understanding how a cell coordinates CDK activation, bistability, and substrate phosphorylation across the cell at mitotic entry ^7, 14, 18, 33, 34^. The predominant view is that the initial mitotic CDK activation is triggered in the cytoplasm at the centrosome, the spindle pole body (SPB) in yeasts, which acts as a mitotic signalling hub controlling the timing of mitosis, from yeast to humans^1–3, 18, 35, 36^. This view has been challenged by *in vitro* experiments suggesting the nucleus controls the timing of mitosis^37, 38^. Differential localization of mitotic regulators, such as Wee1 which is concentrated strongly in the nucleus^39^, also raise the possibility of differences in the bistable responses between the nucleus and cytoplasm^7^. Finally, the timing of CDK substrate phosphorylation has been primarily attributed to the amino acid sequence of the substrates that result in differences in substrate sensitivities, which in combination with a rise in CDK activity order events in the cell cycle^11, 19, 40–42^. However, spatial organization may also influence CDK substrate phosphorylation as a consequence of potentially different CDK activities and substrates being in different cellular locations, which could act in addition to differences in the primary amino acid sequence of substrates.

*Schizosaccharomcyes pombe* (fission yeast) has been an excellent model system to understand core principles in eukaryotic mitotic control^43^. This is partly due to the relative simplicity in its cell cycle control architecture with only one mitotic cyclin-CDK complex, the mitotic B-type cyclin Cdc13, complexed with the Cdk1 homologue, Cdc2^44, 45^. Cell cycle control can be simplified further by engineering a Cdc13-Cdc2 fusion which can drive the entire cell cycle^43^. Furthermore, cell length can be used as a marker for cell cycle position, allowing studies in unperturbed cells^46^. Here we use fission yeast to reveal the underlying spatiotemporal regulatory principles of how CDK orchestrates the onset of mitosis *in vivo*.

### Spatial Differences in CDK Substrate Phosphorylation and Development of Single Cell CDK Sensors

We first asked if there were differences in CDK substrate phosphorylation in different subcellular locations by reanalyzing data from a fission yeast phosphoproteomics experiment, to determine when CDK substrates are phosphorylated during the cell cycle ^11^. We focused on sites which rose in late G2 and during mitosis, using the localization annotation method described in ^47^ to categorize phosphorylation sites based on their cellular compartment^11^. Sites in the nucleus increased before the cytoplasm, cytoplasm periphery, the centrosome/SPB, and the nuclear envelope (Fig 1a, Extended Data Fig. 1a). These results suggest that mitotic CDK activation is not simultaneous across the cell, but rather the initial activation occurs first in the nucleus before activation in other compartments, including the cytoplasm and SPB.

**Figure 1.**
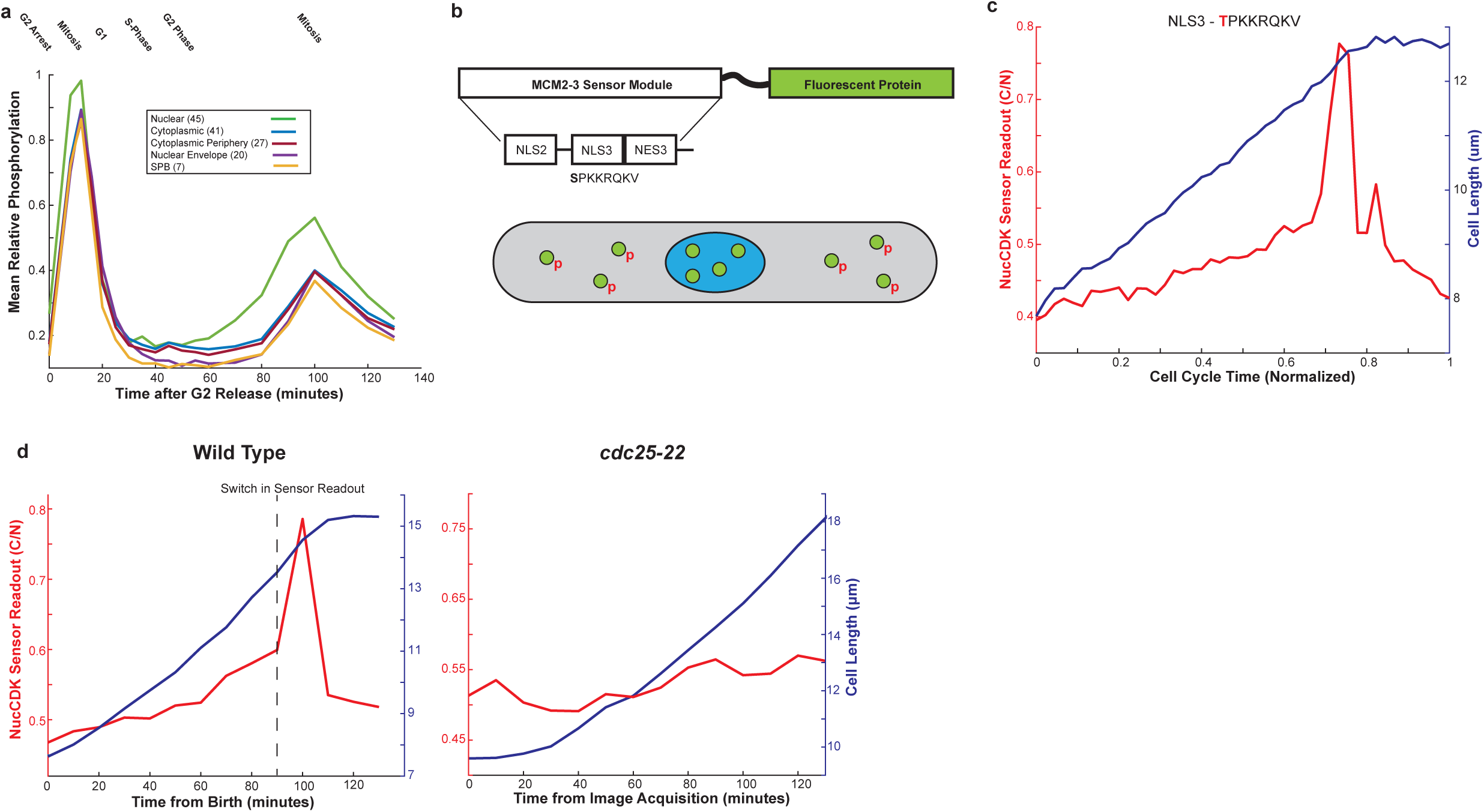
a) Mean phosphorylation of CDK late sites categorized based on annotated localization data. Legend shows number of sites for each spatial compartment in brackets. b) Diagram of minimal phosphorylation module used for construction of NucCDK. NucCDK normally resides in the nucleus (blue) but upon phos-phorylation by CDK, translocates into the cytoplasm (grey). c) Representative profile of NucCDK-mScarletI showing the cytoplasm to nuclear ratio of mean intensity (C/N). Cells were imaged in EMM media with 5 minute time intervals. The alanine mutant of the MCM2-3 module was used as the nuclear mask (MCM2-3 Ala-mNG). d) Representative single-cell traces of NucCDK readout in wild type (left) and *cdc25-22* (right) at 36 degrees, in YE4S media. Images were acquired every 10 minutes. Y-axis for *cdc25-22* is scaled to wild type range.

To investigate this further, we developed new single-cell sensors to assay CDK activity in the nucleus and cytoplasm within unperturbed cells by using cell length as a measure of cell cycle position, combining this with quantitative fluorescence timelapse imaging, simultaneously visualizing cyclin-CDK dynamics.

To assay nuclear CDK activity, we developed a new CDK sensor, NucCDK, which resides in the nucleus but upon phosphorylation by CDK, translocates into the cytoplasm (Fig. 1b, Methods). NucCDK sensor readout increased throughout G2 with a switch to a rapid rise prior to stoppage of cell elongation, a feature of mitotic cells^48^ (Fig. 1c, Extended Data Fig.1d,e). We tested whether this switch occurred at mitotic onset using a temperature sensitive *cdc25-22* mutant that blocks cells in late G2 at 36°C^49^. Unlike wild type cells, there was no rapid switch at 36°C confirming that it was associated with mitotic onset (Fig. 1d). Thus, these switches are a readout of mitotic CDK activation.

### CDK Activation in the Nucleus Drives Mitotic Entry

SynCut3 is a CDK sensor developed in fission yeast for cytoplasmic CDK activity^50^. This sensor is localized in the cytoplasm in interphase but upon phosphorylation by CDK, translocates into the nucleus^50^. We combined this sensor, which we will refer to as CytCDK, together with NucCDK, and performed dual-colour fluorescence imaging to determine the timing of CDK activation simultaneously in both the nucleus and cytoplasm (Fig 2a,b). We used an automated algorithm to identify the point of CDK activation (see Methods). Nuclear activation occurred before cytoplasmic activation, consistent with the phosphoproteomics data (Fig.2c). The time delay between nuclear and cytoplasmic activation was between 5-10 minutes (Fig. 2d, Extended Data Fig 2a). The constancy of the delay between cells suggests that cytoplasmic activation is closely coupled to nuclear activation. We also found there was a delay of 25 minutes between nuclear CDK activation and the stoppage of cell elongation, suggesting CDK is activated earlier in the cell cycle than previously thought (Fig. 2e). A more sensitive version of CytCDK, CytCDK V2, was developed (Methods) that allowed readout of lower levels of CDK activity, and the dual sensor experiments were then repeated. The point representing CDK activation was still the same as when using CytCDK, with a delay of about 5 minutes between nuclear activation and cytoplasmic activation (Fig 2f and g, Extended Data Fig 2b).

**Figure 2.**
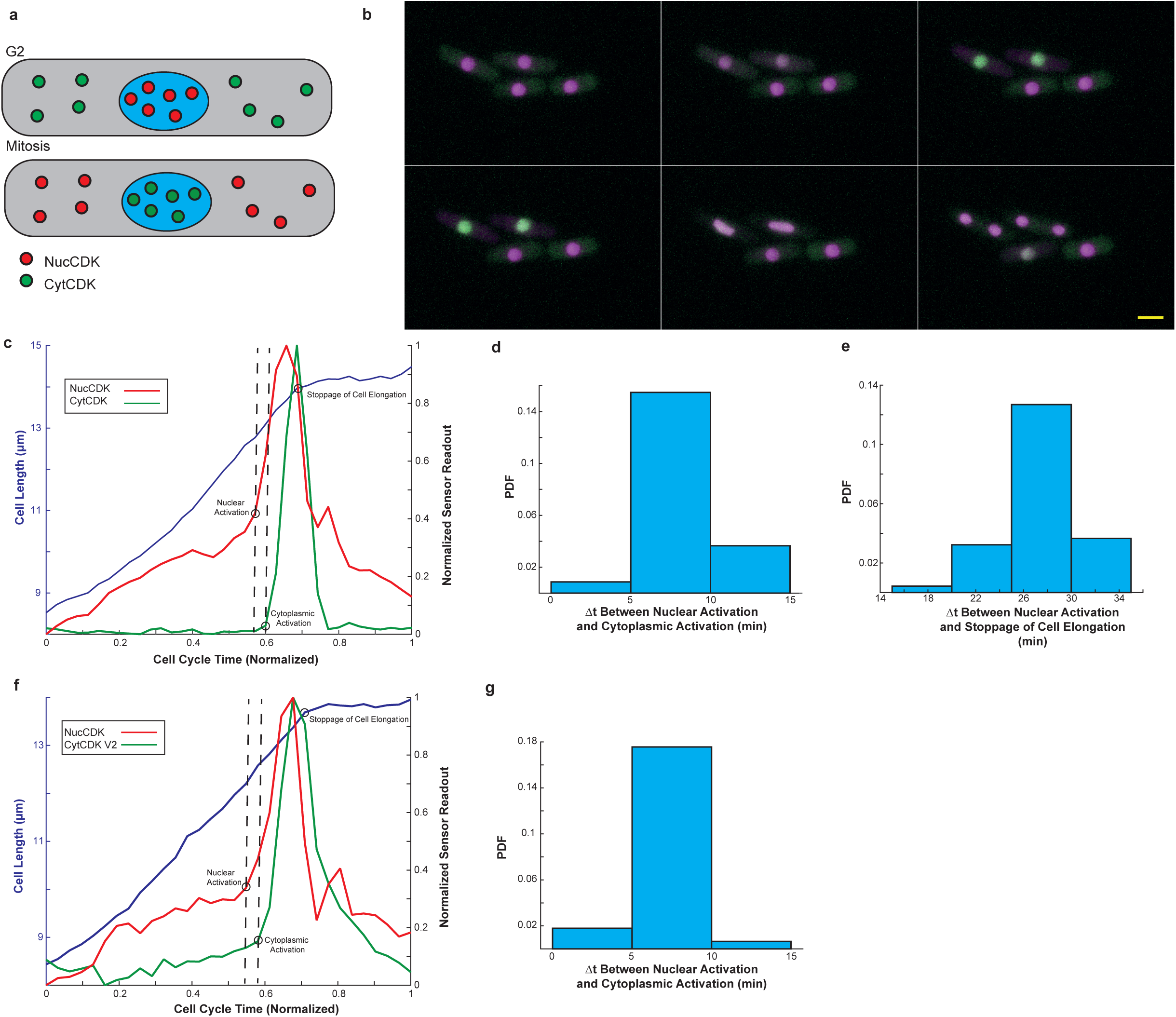
a) Diagram of dual sensor system. b) Montage of dual sensor strain, with NucCDK-mScarletI and CytCDK-mNG, taken every 5 minutes, in YE4S media. Scale bar is 5µm. c) Representative trace of dual sensor results. Black circles indicate points where the rate change algorithm detected a significant change in sensor readout and growth stoppage. Dotted lines shown to represent change points. Sensor readouts were min max normalized. Cell cycle time (normalized) refers to normalizing to the division time for the individual cell. d) Histogram of the probability density function (PDF) showing the time differences from nuclear activation to cytoplasmic activation, in minutes. Bin width was set to time interval used (5 minutes). *n* = 93 cells. e) Histogram showing the time difference from nuclear activation to growth stoppage. Bin width was set to time interval used (5 minutes). *n* = 93 cells. f) Representative trace of dual sensor strain with NucCDK-mScarletI and CytCDK V2-mNG. Experiments were done in YE4S media, with images taken every 5 minutes. g) Histogram of the time differences from nuclear activation to cytoplasmic activation from dual sensor strain with NucCDK-mS-carletI and CytCDK V2-mNG. Bin width was set to time-interval used (5 minutes). *n* = 123 cells.

We conclude that CDK activation occurs first in the nucleus, preceding cytoplasmic activation by 5-10 minutes. This indicates CDK substrate phosphorylation *in vivo* is influenced by where CDK and its substrates are localized in the cell.

### Cdc13-Cdc2 Translocation from the Nucleus is Associated with Cytoplasmic CDK Activation

The translocation of cyclin B1 in human cells at the onset of mitosis is important for mitosis^18, 34^, so we asked if translocation of cyclin-CDK occurs in fission yeast and might couple cytoplasmic activation to nuclear activation. We performed timelapse assays of Cdc13 levels using Cdc13 internally tagged with superfolder GFP (sfGFP) combined with a nuclear mask to segment the nucleus (Methods). We observed a decrease in intensity of Cdc13 in the nucleus just prior to SPB separation, with a corresponding increase in the cytoplasm, indicative of nuclear export at around mitotic onset (Fig 3a, Extended Data Fig 3a,b). This was subsequentially followed by a decrease throughout the cell at mitotic exit after spindle assembly due to Cdc13 degradation at mitotic exit (Fig 3a, Extended Data Fig 3a,b). For further experiments, we used the mean of the top 15% of pixels as an estimate of nuclear intensity since nuclear area is 15% of cell area ^51^. This gave similar results to using a nuclear mask (Extended Data Fig. 3c), thereby avoiding use of a nuclear mask to minimize light exposure. Timelapse imaging of Cdc2-mNeonGreen (mNG) indicated a similar nuclear export happening prior to spindle assembly, which was followed by further export after Cdc13 degradation (Fig. 3b, Extended Data Fig.3d). We performed dual colour imaging of Cdc13-sfGFP with NucCDK and CytCDK to identify when Cdc13 export occurs in relation to nuclear and cytoplasmic activation. We found nuclear activation occurred about 5 minutes before export of Cdc13 while cytoplasmic activation occurred concurrently with export (Fig. 3c,d,e). A similar 5-minute difference was observed for Cdc2-mNG with respect to nuclear activation, suggesting that Cdc13 and Cdc2 translocate as a complex (Extended Data Fig. 3e). When we calculated the nuclear and cytoplasmic mean Cdc13-sfGFP intensities at nuclear and cytoplasmic activation we found the nucleus had a higher value, indicating differences in the cyclin threshold for CDK activation between the nucleus and cytoplasm (Fig.3f, Extended Data Fig 3f). This suggests that CDK is first activated in the nucleus where there is a higher cyclin threshold, and this is followed by a fraction of that cyclin-CDK translocating to the cytoplasm, which overcomes the cyclin threshold in the cytoplasm for cytoplasmic CDK activation.

**Figure 3.**
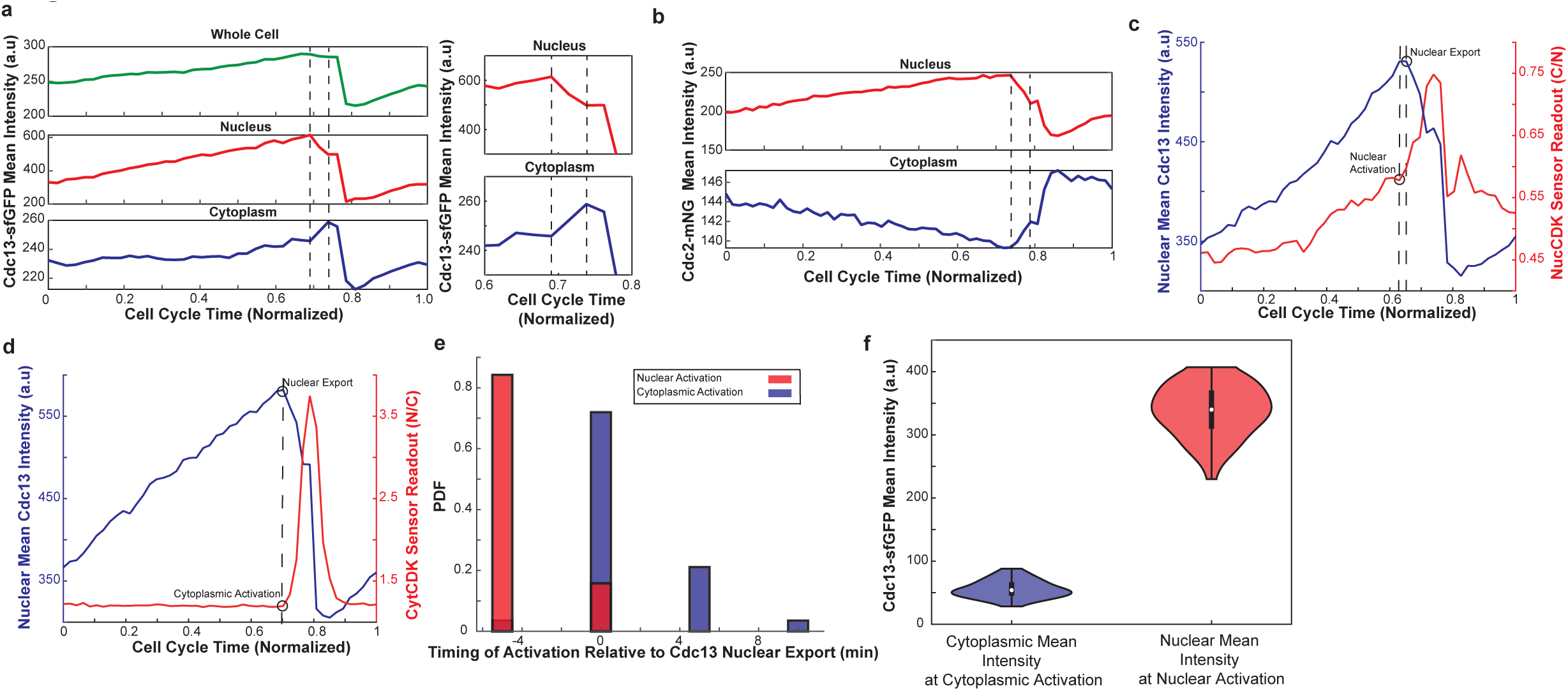
a) Left) Representative trace showing mean intensities of whole cell, nucleus, and cytoplasm, using MCM2-3-Ala -mScarletI as nuclear mask. Dotted lines mark beginning and end of nuclear export. Right) Zoomed in trace to better visualize translocation of Cdc13-sfGFP. Cells were imaged using EMM media, using 5 minute intervals. b) Representative time trace of Cdc2-mNG mean intensity in nucleus and cytoplasm calculated using top15% method. Dotting lines mark beginning and end of nuclear export. Cells were imaged using EMM media, with 5 minute intervals. c) Representative trace of NucCDK-mScarletI and Cdc13-sfGFP showing nuclear activation with respect to nuclear export. Cells were imaged using EMM media, with 5 minute intervals. d) Representative trace of CytCDK-mScarletI and Cdc13-sfGFP showing cytoplasmic activation with respect to nuclear export. Cells were imaged using EMM media, with 5 minute intervals. e) Histogram of nuclear activation (*n* = 76 cells) and cytoplasmic activation (*n* = 57 cells) with respect Cdc13-sfGFP nuclear export in minutes. Red represents data from NucCDK-mScarletI and Cdc13-sfGFP and blue represents data from CytCDK-mScarletI and Cdc13-sfGFP. f) Violin plots of nuclear mean intensity and cytoplasmic mean intensity after background subtraction, at nuclear (*n* = 76 cells) and cytoplasmic CDK activation (*n* = 57 cells), respectively. Mean background values of the nucleus and cytoplasm were calculated as the mean of the median values shown in Extended Data Fig. 3f and subtraced from each value.

### CDKY15 Feedback Loops Provide Greater Stability in Oscillations in the Nucleus

Bistability (multiple stable steady states of system) and hysteresis (state of system dependent on its history) in Cdk1 activity has been shown *in vitro* with *Xenopus* egg extracts using steady state measurements of Cdk1 activity to non-degradable cyclin^6, 8^. However, hysteresis in CDK activity has not been directly tested *in vivo,* and potential spatial differences have not been investigated in any system. Therefore, we investigated the different hysteretic/bistable responses between the nucleus and cytoplasm, and whether hysteresis may permit translocation of Cdc13-Cdc2 to the cytoplasm without collapse of the mitotic state in the nucleus. To address this, we determined the dynamic single-cell phase plots in the nucleus and cytoplasm *in vivo* as they represent the naturally occurring phase orbits occurring in the dynamical system^5^. The phase plots allow visualisation of the different states of CDK activity for different concentrations of cyclin^4–6^. The phase plots using data from Fig. 3c-f indicated a significant difference in the phase orbits between the nucleus and cytoplasm (Extended Data Fig. 3g).

This was confirmed using the Cdc13-Cdc2 fusion protein tagged with mNG, which confines analysis to a single cyclin-CDK complex^43^, along with NucCDK and CytCDK V2. The dynamics of Cdc13-Cdc2-mNG mimicked uncomplexed Cdc13, including nuclear export, along with similar sensor readout profiles (Extended Data Fig.4 a-d). The nuclear and cytoplasmic phase orbits were plotted on the same phase space, after min max normalization of sensor readouts (Fig. 4a). Strikingly, the nuclear phase orbit cycled around a much larger region of phase space than the cytoplasm(Fig 4a). A high amount of Cdc13-Cdc2 was able to accumulate in the nucleus before CDK activation to the high activity mitotic state, with little increase in activity prior to that point. Once in the mitotic state, the nucleus was able to tolerate decreases in Cdc13-Cdc2 without collapsing back to the low activity state. Furthermore, for the same mean intensity of Cdc13-Cdc2, the sensor readout was either low or high, depending on what state the nucleus was in, consistent with bistability and hysteresis. This is in contrast to the cytoplasm in which very little Cdc13-Cdc2 accumulation was needed before CDK activation, and very little decrease in Cdc13-Cdc2 resulted in mitotic state collapse (Fig. 4a). Overall, the phase orbit in the cytoplasm was much more narrow compared to the one in the nucleus. This suggests that the nucleus is much more stable with respect to cyclin-CDK fluctuations compared to the cytoplasm. The individual nuclear phase plots extended and shrunk significantly horizontally compared to the cytoplasm, also suggesting less sensitivity to cyclin-CDK concentrations (Fig 4a, bottom).

**Figure 4.**
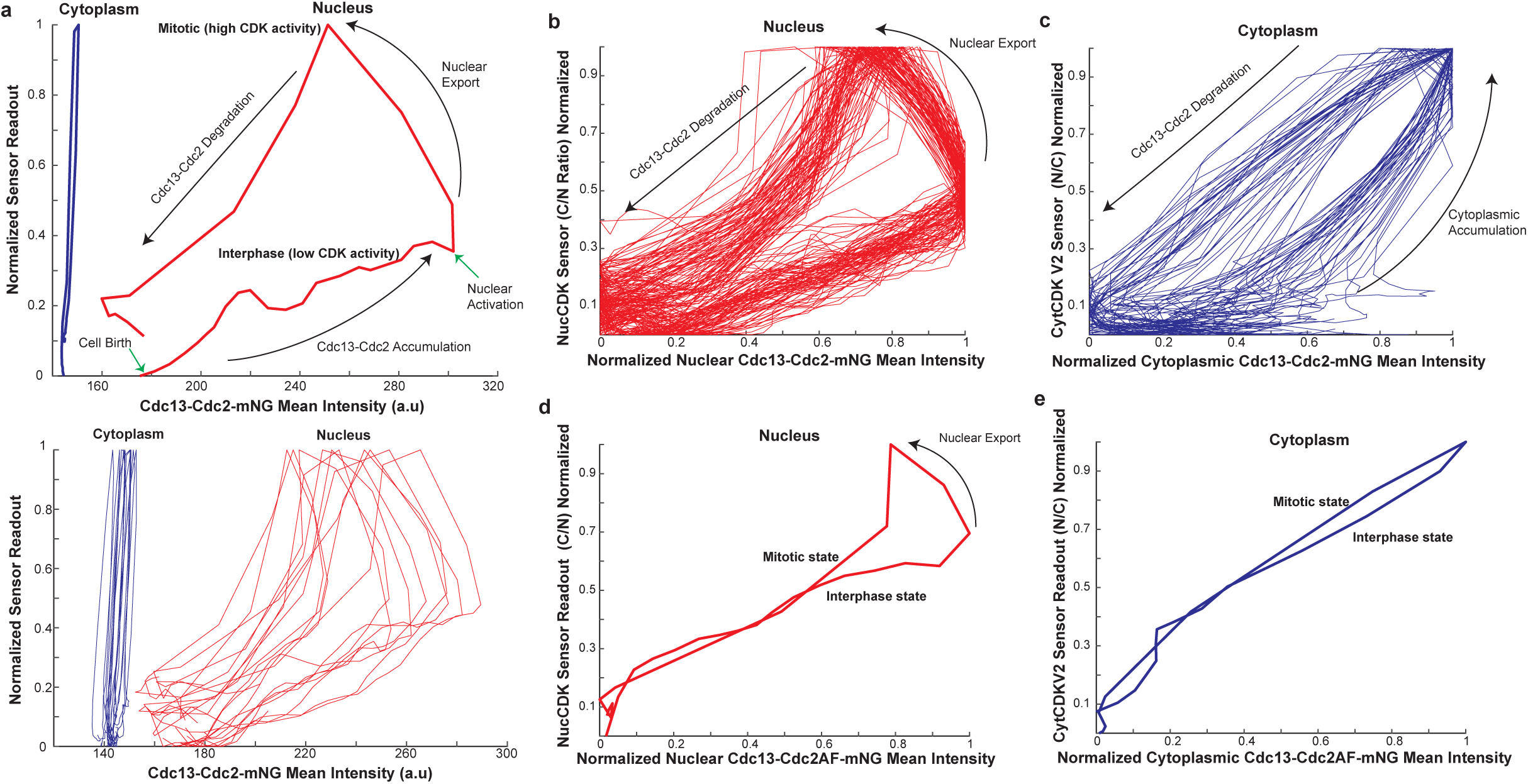
a) Top: Representative phase plot of the nuclear phase orbit (Cdc13-Cdc2-mNG nuclear mean intensity and NucCDK-mScarletI), and cytoplasmic phase orbit (Cdc13-Cdc2-mNG cytoplasmic mean intensity and CytCDK V2-mScarletI), plotted on the same phase space. Sensor readouts were min max normalized for comparison. Bottom: 10 representative traces plotted for the nucleus and cytoplasm each. b) Min max normalized phase plots of the nucleus in Cdc13-Cdc2-mNG background. *n* = 89 cells c) Min max normalized phase plots of the cytoplasm in Cdc13-Cdc2-mNG background. *n* = 37 cells d) Representative min max normalized phase plot of the nucleus in the Cdc13-Cdc2AF-mNG background. e) Represenative min max normalized phase plot of the cytoplasm in the Cdc13-Cdc2AF-mNG background.

When we plotted the single-cell phase plots of the nucleus after min max normalization of Cdc13-Cdc2 mean intensities, we noticed a considerable gap between the low activity and high activity states, and that after nuclear export, the system was not prone to collapse back to the low activity state(Fig. 4b). Furthermore, the orbits were very similar in the different single cells, suggesting that the CDK regulatory system oscillates consistently and is robust to biological noise. In contrast, the normalized cytoplasm phase plots showed that the gaps between the high activity and low activity states for the same Cdc13-Cdc2-mNG intensity, were less pronounced and the cytoplasm was more sensitive to Cdc13-Cdc2 accumulation(Fig. 4c). The orbits were also similar in different cells.

We repeated the experiments using the Cdc2T14AY15F (Cdc2AF) mutant fused to Cdc13 (which is not viable unless fused to Cdc13), Cdc13-Cdc2AF-mNG, which abolishes the CDKY15 feedback loops to test if CDKY15 feedback loops are responsible for the stability and the distinction of the low and the high states in the nucleus and cytoplasm^5^. Nuclear export appeared less switch-like, as the drop in nuclear intensity and accumulation in the cytoplasm was less pronounced, suggesting the export is influenced by the altered CDK activity regulation (Extended Data Fig. 4e,f). The sensors displayed a linear increase in sensor readout (Extended Data Fig. 4g,h). Both the phase diagrams of the nucleus and cytoplasm exhibited almost a complete collapse of their orbits (Fig. 4d,e, Extended Data Fig. 4i,j). Thus, CDKY15 feedback loops provide hysteretic responses in both the nucleus and cytoplasm. In the nucleus, when export occurs, the system is subsequently prone to collapse, indicating that in the Cdc13-Cdc2AF, translocation comes with the risk of the nucleus collapsing back to the interphase state (Fig. 4d). We conclude that CDKY15 feedback loops provide a strong hysteretic response in the nucleus, promoting translocation of cyclin-CDK from the nucleus to the cytoplasm, and a much weaker response in the cytoplasm.

### The Role of Cyclin-CDK at the Yeast Centrosome

These results differ from the predominant view that mitotic CDK activation is triggered first at the centrosome/SPB thereby dictating the timing of mitotic entry^1, 3, 36^. Therefore, we investigated what role cyclin-CDK at the SPB might play at the onset of mitosis. In fission yeast, CDK activation at the SPB has been shown to act through a polo kinase (Plo1) mediated feedback loop, dependent on Plo1 association with the SPB scaffold protein, Cut12(Fig. 5a)^1, 52–55^. The mutant *cut12.s11* promotes earlier recruitment of Plo1 to the SPB and global Plo1 activation^53, 54^ so we determined if *cut12.s11* advanced the timing of nuclear CDK activation (Fig. 5b). Cell length at nuclear CDK activation assayed using NucCDK (Fig. 5c, left), and cell length at cell division (Fig. 5c right), were found to be the same in both *cut12.s11* and wild type cells. This indicates that promoting premature CDK activity at the SPB has no significant impact on either nuclear CDK activation or the timing of mitosis. This is in line with earlier reports that promoting SPB feedback in a normal cellular context does not have an impact on mitotic timing^52^.

**Figure 5.**
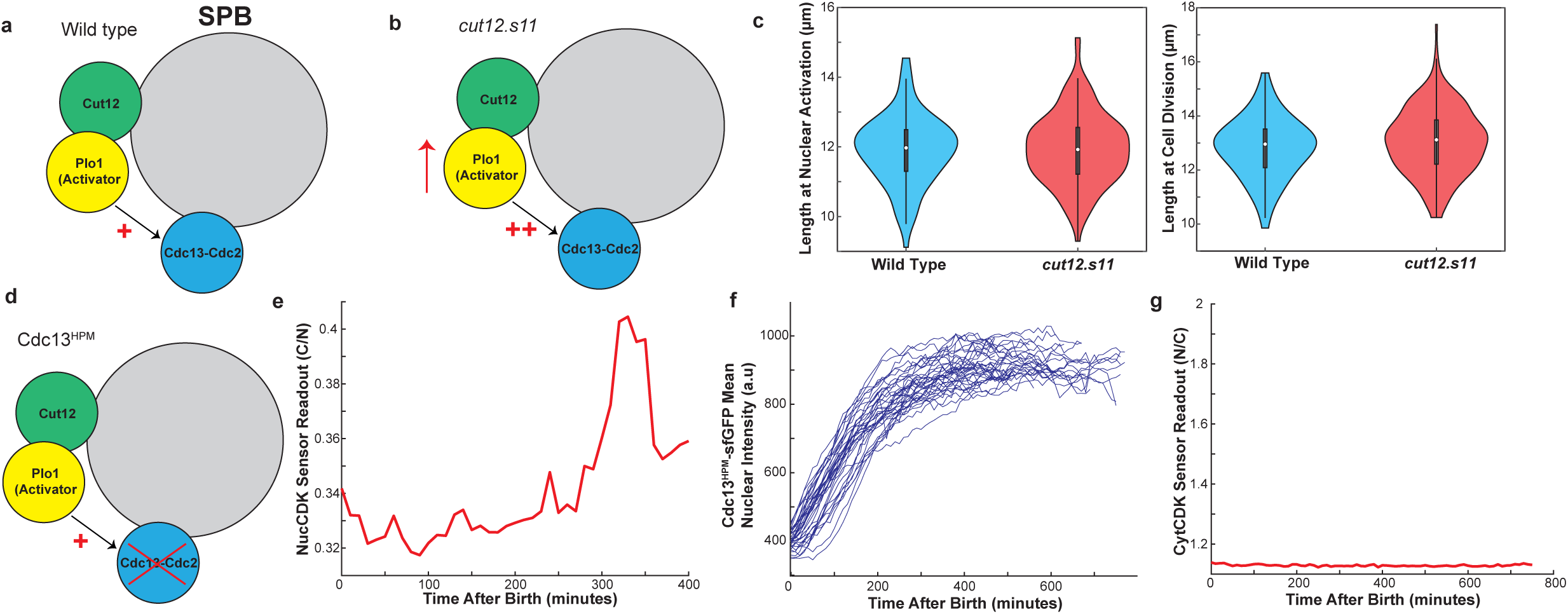
a) Diagram of the SPB in a wild type scenario with Plo1 activating Cdc13-Cdc2 at the SPB. b) Diagram of the SPB in a *cut12.s11* mutant showing increased recruitment of Plo1 to the SPB through Cut12, thereby promoting activation of Cdc13-Cdc2 at the SPB. b) Left) Violin plot comparison between wild type cells (*n* = 96 cells) with NucCDK-mScarletI and *cut12.s11* cells (*n* = 195 cells) with NucCDK-mScarletI showing length at nuclear activation. Right) Violin pot comparison of wild type cells (*n* = 96 cells) and *cut12.s11* cells (*n* = 195 cells) of length at cell division. d) Diagram of the SPB in the Cdc13^HPM^ background, where Cdc13 SPB localization is abolished. e) Representative time trace of NucCDK-mScarletI in Cdc13^HPM^-sfGFP background. f) Traces of mean nuclear Cdc13^HPM^-sfGFP intensity. *n* = 36 cells. g) Representative time trace of CytCDK-mS-carletI in Cdc13^HPM^-sfGFP background. Y-axis of CytCDK sensor readout was scaled based on the minimum and maximum value of CytCDK in Cdc13^WT^.

We next investigated the role of cyclin-CDK localization at the SPB for mitotic control. For this, we used the hydrophobic patch mutant (hpm) of Cdc13 (Cdc13^HPM^), which does not localize to the SPB during G2^47^ (Fig. 5d), a property conserved with mammalian cyclin B1^HPM^ which also does not localize to the centrosome^47^. Cells with Cdc13^HPM^ alone do not display signs of mitotic entry, which is partly rescued by tethering Cdc13^HPM^ back to the SPB^56^. We performed timelapse microscopy of Cdc13^HPM^-sfGFP and wild type Cdc13 (Cdc13^WT^)-sfGFP, using the CDK sensors (see Methods) and with other cyclins deleted. Cdc13^WT^-sfGFP displayed normal sensor readouts and Cdc13 dynamics (Extended data Fig.5a,b,c). NucCDK in the Cdc13^HPM^-sfGFP mutant indicated that activation in the nucleus still occurred, even though Cdc13^HPM^-sfGFP did not localize to the SPB (Fig. 5e, Extended Data Fig. 5d,e). However, strikingly this nuclear CDK activation was not followed by a drop in nuclear levels indicating export from the nucleus did not take place (Fig. 5f).

We repeated the timelapse experiments with Cdc13^HPM^-sfGFP and CytCDK and found cytoplasmic CDK activation was also absent (Fig. 5g, Extended Data Fig. 5f). Therefore, while nuclear activation still occurs, cytoplasmic CDK activation is significantly impaired as a likely consequence of Cdc13-Cdc2 not being exported from the nucleus.

We propose that Cdc13 localization to the SPB is not required for CDK activation in the nucleus, but is required for the translocation of Cdc13-Cdc2 to the cytoplasm and the subsequent activation of cytoplasmic CDK. These results are consistent with previous phosphoproteomics data^47^, which showed phosphorylation of nuclear substrates in Cdc13^HPM^ was similar to Cdc13^WT^, whilst phosphorylation of SPB and cytoplasmic substrates was significantly impaired.

## Discussion

The onset of mitosis is a critical transition in the eukaryotic cell cycle but the core principles of how CDK regulates it *in vivo* are not well understood. Here we have revealed the underlying spatiotemporal regulatory framework in fission yeast as to how CDK orchestrates the onset of mitosis *in vivo* (Fig. 6.a,b). This work has shown that a full understanding of CDK control over mitosis needs to take account of cellular compartmentation, requires precise measurements of the levels of CDK and its regulators, the sensing of *in vivo* CDK activity and the generation of phase plots, carried out using single unperturbed cells. This is possible in model eukaryotes like fission yeast which can be precisely manipulated genetically and in which the CDK regulatory circuits can be simplified to accommodate the redundancy, plasticity and overlapping functions of CDK regulatory components, all of which make interpretation of experiments difficult^14, 43^.

**Figure 6.**
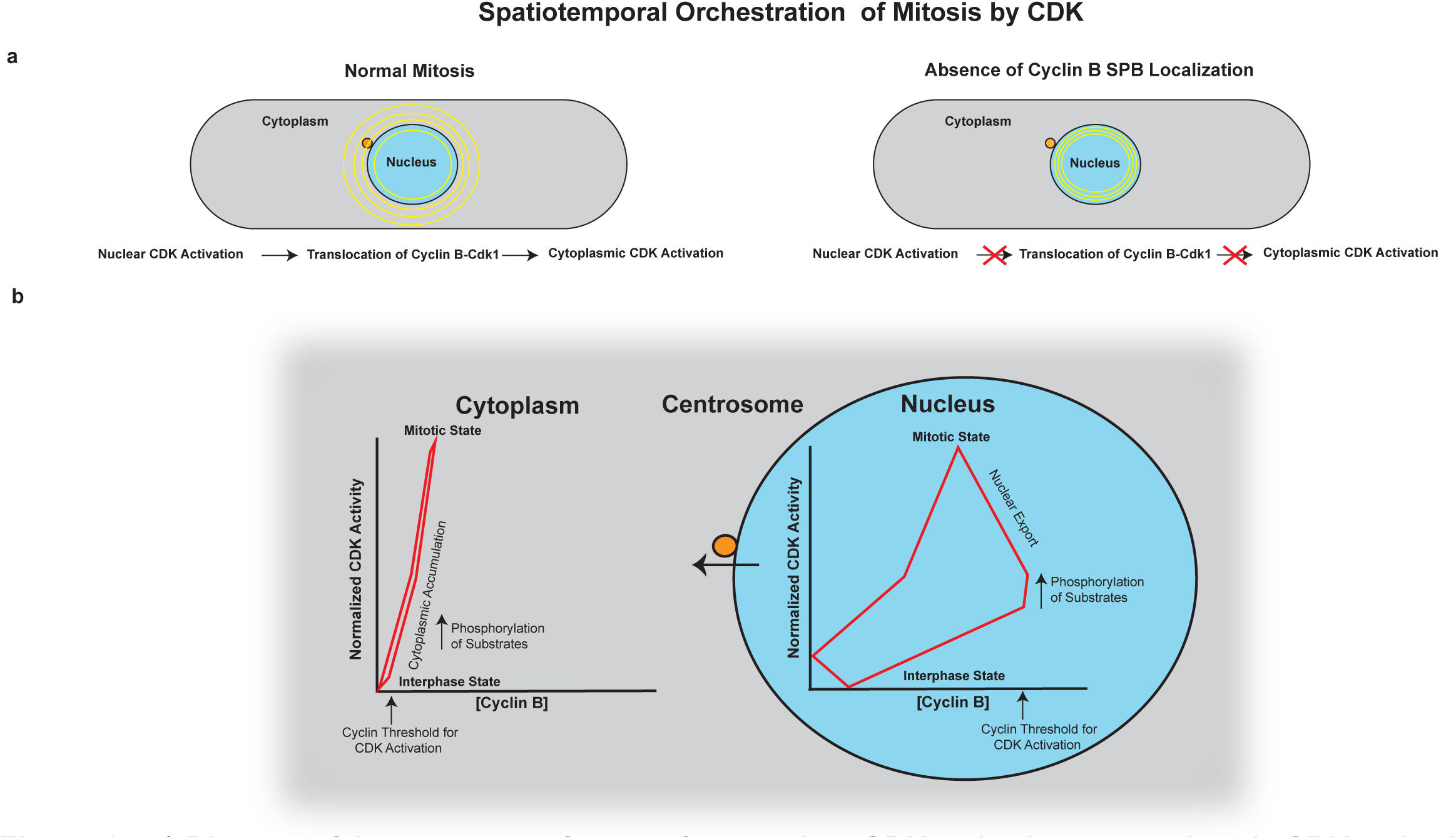
**a)** Diagram of the sequence of events from nuclear CDK activation to cytoplasmic CDK activation and the role of SPB localization of cyclin B. b) Diagram of the phase orbits in the nucleus and cytoplasm. During G2, cyclin B-CDK concentration increases in the nucleus but CDK activity only increases gradually, with a high cyclin B threshold for activation. In late G2, there is a rapid, switch-like rise in CDK activity which is dependent on CDKY15 feedback loops, that also increases the phosphorylation of substrates localized in the nucleus. After this point, cyclin B-CDK translocates from the nucleus to the cytoplasm for signal propagation, but high CDK activity is still maintained in the nucleus due to CDKY15 feedback loops. Thus, the oscillatory orbit is very stable with respect to cyclin B-CDK concentration in the nucleus. In contrast, the accumulation of cyclin B-CDK in the cytoplasm immediately promotes CDK activity through CDKY15 feedback loops, albeit to a significantly lesser degree than in the nucleus. This results in increased phosphorylation of cytoplasmic substrates. There is also a lower cyclin B threshold for activation in the cytoplasm than in the nucleus, allowing cytoplasmic CDK to rapidly respond to the import of cyclin B-CDK from the nucleus. However, within the cytoplasm the oscillatory orbit is less stable with respect to cyclin B-CDK concentration.

Our *in vivo* work has shown that CDK is first activated in the nucleus, and dictates the timing of mitotic onset and cell cycle progression. This is in contrast to the prevailing model that CDK is activated in the cytoplasm first, specifically the centrosome/SPB, which is thought to be a mitotic signalling hub controlling the timing of mitotic onset^1–3, 36^. We found that promoting activity at the SPB had no effect on the timing of nuclear activation nor timing of mitosis. However, in the absence of cyclin B-CDK localization to the SPB, CDK activation still occurred in the nucleus but not in the cytoplasm. This effectively results in spatial uncoupling of mitotic onset with one compartment entering the mitotic state (nucleus) while the other remains in a G2-like state (cytoplasm). We propose this is due to the absence of signal propagation, in the form of nuclear export of cyclin B-CDK to the cytoplasm. This stresses the importance of signal propagation for spatially coherent mitotic onset and suggests that the SPB does act as a signalling hub, but as a signal relaying hub as opposed to one controlling the timing of mitotic onset.

We also generated novel *in vivo* single-cell phase plots of CDK activity versus concentrations of cyclin B and of cyclin B-CDK complexes simultaneously in both the nucleus and cytoplasm. These showed that the CDK oscillator operates in very different domains within the nucleus and cytoplasm. Within the nucleus, there is a strong bistable/hysteretic response, which can buffer against fluctuations in cyclin B-CDK concentrations in both interphase and mitosis, resulting in stable oscillations in CDK activity. In contrast, within the cytoplasm the response was much weaker. The strong bistable response and higher cyclin B threshold for CDK activation in the nucleus reduces susceptibility to ‘noise’ and to the risk of slipping in and out of mitosis helping maintain genomic stability^5, 32^. This regulation could be due to the higher concentrations of CDK regulators in the nucleus such as Wee1 and Cdc25^7, 39, 51^. In contrast, the less well stabilised CDK circuit and lower cyclin B threshold in the cytoplasm allows it to rapidly respond to the translocation of cyclin B - CDK from the nucleus to the cytoplasm. Therefore, the cytoplasm may exist in a more fluid state where the distinction between interphase and mitosis is less clear. The translocation of cyclin B-CDK would link the different oscillatory orbits between the nucleus and cytoplasm leading to coherent mitotic entry, resulting in the nucleus enforcing control in the cytoplasm^7^. Theoretical modelling of the metazoan mitotic network^7^ has proposed similar differences between the nucleus and cytoplasm in bistable responses and cyclin B thresholds to those which have been shown here experimentally. Bistability is a common characteristic of biological signalling networks and cell transitions^6, 57, 58^ and we have shown that the responses can be very different between cellular compartments.

Modelling of the mammalian mitotic network has suggested that nuclear envelope breakdown (NEBD) creates a potential “stress” at mitotic entry^59^, due to dispersion of cyclin B1-Cdk1 and Greatwall kinase resulting in mitotic collapse – a collapse that can be prevented by Cdk1Y15 feedback loops which retain high CDK activity^59^. The nuclear export of cyclin B-CDK observed here in fission yeast which has a closed mitosis, can potentially lead to a similar stress in the nucleus that is also prevented by CDKY15 feedback loops which help retain high CDK activity. This would ensure nuclear mitotic events such as chromosome condensation and mitotic spindle formation can continue to proceed. In contrast, the cytoplasm is ill-suited for propagating CDK activity because of its weaker bistable response. We propose that the strong bistable response in the nucleus allows propagation of CDK activity from the nucleus to the cytoplasm — a potential mitotic stress — without collapse of nuclear CDK activity. Although the fission yeast cyclin B-CDKAF is viable, mitosis is faulty. It is sluggish with defective behaviours, exhibits less well-regulated timing of mitotic entry, and generates a higher frequency of inviable cells^43, 50^.

The fission yeast CDK regulatory framework can be expected to be relevant to other eukaryotes. The *in vitro Xenopus* extract studies proposing that the nucleus is the “pacemaker” for the cell cycle^37, 38^ suggest that our observations are not simply due to differences between a closed (yeast) vs open mitosis. In human cells, the mechanism behind cyclin B1 nuclear translocation prior to NEBD is thought to be due to either Cdk1 phosphorylation of transport machinery or through a spatial positive feedback mechanism based on cyclin B1 autophosphorylation whereby the nucleus promotes the translocation of cyclin B1^18, 34^. This translocation is important for triggering NEBD^19^. Furthermore, it has been proposed that robust activation of cyclin B1-Cdk1 occurs in the nucleus^35, 60^. Therefore, one possibility is that activation of cyclin B1-Cdk1 may also occur first in the nucleus, with translocation of cyclin B1-Cdk1 into the nucleus resulting in further activation. During this time, some active cyclin B1-Cdk1 from the nucleus may diffuse into the cytoplasm increasing cytoplasmic activity, but upon NEBD cyclin B1-Cdk1 disperses, leading to robust cytoplasmic activation. In addition, cyclin A2, which is mainly nuclear^15, 33^, has been proposed to inactivate the nuclear-enriched Wee1^39^ to lower the activation threshold of cyclin B1-Cdk1^15, 16^, which could initiate nuclear cyclin B1-Cdk1 activation and translocation into the nucleus. In fission yeast, the translocation of Cdc13-Cdc2 from the nucleus to the cytoplasm could be analogous in function to translocation of cyclin B1-Cdk1 triggering NEBD, resulting in similar robust cytoplasmic CDK activation. The general underlying principle is CDK regulation of its own translocation. This is demonstrated here with the Cdc2AF mutant perturbing CDK activation that resulted in less abrupt translocation, which is similar to what has been shown with human cyclin B1 and Cdk1AF^34^. Therefore, although the precise mechanisms may have specific differences, the general concepts of the nucleus acting as the mitotic pacemaker and translocation being important for propagating CDK activity appear to be conserved regardless of differences between open and closed mitosis^18, 37, 38^.

We have also shown that in addition to the amino acid sequence of substrates contributing to their phosphorylation profile^40–42^, the *in vivo* localization of substrates also plays a key role due to the different CDK activation responses between the nucleus and cytoplasm. Within a spatial compartment, the rise in CDK activity along with substrate phosphorylation thresholds, will trigger sequential cell cycle events. Some of these events, such as the export of Cdc13-Cdc2 into the cytoplasm, affect CDK activity in other compartments, thereby triggering substrate phosphorylation. Therefore, the timing of CDK substrate phosphorylation is determined by an interplay between primary amino acid sequence, CDK and substrate localization, and specific events that occur during the cell cycle.

Why does the initial mitotic CDK activation and strong bistable/hysterisis response occur in the nucleus? A major vulnerability of the mitotic cell cycle is the risk of premature CDK activation and mitotic onset when DNA is incompletely replicated or has unrepaired damage^61^. Having the CDK regulatory system within the nucleus, positions the bistable circuit close to DNA, so incomplete DNA replication and DNA damage can be readily integrated with CDK activity control^62, 63^. Such a tight association between DNA and CDK activation would help ensure genome integrity. It also allows genome content (ploidy) to be more easily monitored, which is necessary given cell size at mitosis increases with ploidy^50, 64^. Thus, DNA may serve as a platform for CDK activity control and the initiation of mitotic entry^50, 62^.

The principles we have developed here will be a useful guide for future experiments in the more complex metazoan cells investigating how CDK brings about cell cycle progression. Furthermore, the signaling network in the CDK oscillator system has been shown to have similar features to other biological oscillators, such as bistability and positive feedback loops^65^. Therefore, our work here will also be useful for understanding how other biological oscillators operate across space and time *in vivo*.

## Methods

### Development of NucCDK and CytCDK sensors

All the sensors were expressed under the low expression constitutive calmodulin (cam1) promoter^66^. The sensors were tagged with either mNeonGreen or mScarletI^67, 68^. NucCDK was derived from a minimal budding yeast CDK-dependent transport module, that was shown to be exported out of the nucleus after phosphorylation by Cdc28 (budding yeast Cdk1) (Fig.1b)^69^. The MCM2-3 sensor module displayed a readout patten that was very similar to that of early CDK substrates that rise at the onset of S-phase, consistent with observations in budding yeast (Extended Data Fig. 1b)^11, 69^. Although this version of the sensor is useful for the G1/S transition given its high sensitivity to CDK activity^42^, we aimed to reduce its sensitivity so it can read out activity at G2/M. The module contains a full CDK consensus site beginning with a serine, next to the Mcm3 NLS. Mutation of this residue to alanine (MCM2-3 Ala) drastically reduced the readout of the sensor and rendered it prominently nuclear, albeit with a very modest increase after stoppage of cell elongation (Extended Data Fig. 1c). Therefore, we hypothesized that mutating this serine to a threonine could alter the sensitivity of the sensor as it has been shown that CDK has a much greater specificity towards serine residues than threonine residues, and furthermore, that threonines are preferentially dephosphorylated by PP2A^41, 70^. As shown in Extended Data Fig.1d, this mutation drastically altered the profile of the sensor, and gave a readout of CDK activity throughout the cell cycle. We refer to this sensor as NucCDK. The profile was very similar to a recent CDK Forster resonance energy transfer (FRET) sensor tagged with a nuclear localization sequence, developed in fission yeast, supporting that NucCDK is biased towards sensing nuclear CDK activity^71^. We did not notice a significant difference in the pattern when using a nuclear mask for determining the cytoplasmic to nuclear ratio in mean intensities, as opposed to using the mean of top 15% of pixels and bottom 85% of pixels as estimates for nuclear and cytoplasmic mean intensities, respectively, as has been previously used in fission yeast (Extended Data Fig. 1e)^51^. Unless stated that a nuclear mask was used, the top 15% and bottom 85% method was used for sensor readout measurements.

Syncut3 (CytCDK) translocation into the nucleus is dependent on phosphorylation of T19 by CDK^50^. CytCDK V2 was developed in a similar manner to NucCDK except T19 was mutated to a serine, for increased sensitivity to CDK activity.

### Phosphoproteomics Timecourse Analysis

The timecourse analyzed for Fig. 1a was from the Cdc13-Cdc2 fusion experiment where Cig1, Cig2, Puc1 had been deleted (ΔCCP), grown in EMM4S + SILAC (stable isotope labeling with amino acids in cell culture) media at 32° ^11^. Localization annotation for CDK sites was implemented as previously described in ^47^. In brief, the order of determining the localization of a protein by priority based on existing literature was as follows: 1) Direct visualization of fluorescently tagged proteins labelled at the endogenous locus. 2) Fluorescently tagged proteins from exogenous nmt41 or nmt81 promoters. 3)Indirect methods suchs as ChIP assays, fractionated western blotting, and immunoprecipation experiments. Only sites annotated as being late sites as well as belonging to solely one spatial compartment were used for analysis.

### Growth and Strain Construction

Growth of strains and cell cultures, and media was done as described in^72^. Strains were constructed either by transformation or genetic crossing, as described in^72^. All experiments were done in exponentially growing cells of 2.5-4.0 x 10^6^ cells/ml. Insertions were checked with colony PCR and sequencing in the case of deletions. All strains and plasmids used are listed in Supplementary Table 1. Edinburgh minimal media (EMM) powder (MP Biomedicals) with ammonium chloride and dextrose was used, supplemented with adenine at 0.15g/L final concentrations, where required. Where yeast extract with adenine, leucine, histidine, and uridine (YE4S) was used, supplements were added at 0.15g/L final concentrations.

The Cdc13 strain internally tagged with sfGFP is described in^73^.

### Microscopy

Except for data in Fig. 4, microscopy was done on 1% agarose pads, using GeneFrames, to prevent drying of pad during acquisition^74^ (125ul, Thermofisher AB-0578). N-propyl gallate (Sigma, 02370) was added to the agarose pads at 0.1mM final concentration to prevent photobleaching^75^.

A lml culture of cells were spun down once at 2000RPM for 30s, to minimize stress. Approximately 50ul of supernatant was left and mixed. 2ul of cell culture was spread across agarose pad and left to dry for a few minutes. A coverslip (Epredia, 22x32mm, 1.5) was placed onto the pad, but only pressure along the gene frame was applied to seal up the frame. Nail polish remover was added along the frame to ensure that the coverslip and frame were sealed.

Imaging was done on a Nikon Ti2 inverted microscope equipped with a perfect focus system (PFS), a Okolab environmental chamber, and a Prime BSI sCMOS camera (Photometrics). A Nikon 100x Plan Apo oil immersion lens with a NA of 1.45 was used for imaging. Image acquisition was controlled using µManager V2.0 software^76^. 2x2 pixel binning was done to improve fluorescence signal detection with a resulting pixel size of 130nm x 130nm. Unless stated otherwise, imaging was done at 25°. Except for data shown in Extended Data Fig 2a, where 5 z-slices were collected, 7 slices were collected with 0.5um step sizes. Fluorescence excitation was done using a SpectraX LED light engine (Lumencor). Imaging for mNG-tagged proteins was done with the YFP channel (both in excitation and emission) to minimize the influence of cell autofluorescence^77^. We did not notice any issues with growth or cell length at division under the imaging conditions used.

### Cdc13^HPM^ experiments

Cells contained a copy of Cdc13^WT^or Cdc13^HPM^ expressed under the native Cdc13 promoter and a second copy of Cdc13^WT^, expressed under the thiamine-repressible nmt41 promoter^47^. Cells were initially grown in EMM media without thiamine but for experiments, cells were washed once in EMM + thiamine and placed on a 1% EMM + thiamine agarose pad. Thiamine was added to EMM at a final concentration of 30uM. Only cells that were born around the beginning of acquisition and blocked at the end of acquisition were considered for analysis. The total acquisition was 80 time points with time intervals of 10 minutes, for around 13 hours in total in acquisition.

### Data and Image Analysis

Unless state otherwise all analysis was done using custom scripts written in Matlab R2022a. For processing and analysis of fluorescence z-stacks, maximum intensity projections were used using Fiji^78^. Cells were segmented using a custom algorithm described in^56^. Nuclear masks were obtained using Fji, using MCM2-3 Ala to identify the nucleus. Cells were tracked using LineageMapper^79^. With the exception of cases where mutants that block cells in G2 were used, only cells that were born and divided within the timelapse acquisition period, were used for analysis.

For histograms, the left edges of bins are inclusive, while the right edges are not inclusive except for the last bin. Violin plots display distributions as kernel density estimates. The median value is plotted as a white circle. The black rectangle represents the interquartile range while the black solid line represents 1.5x interquartile range.

### Rate Change analysis

Sensor traces were lightly smoothed using the Matlab function, “smoothdata”, and the algorithm, “sgolay”, to assist in identifying change points. However, plots in figures represent the raw data. To confine analysis to the period in G2 where the rate change in sensor readout occurs, the trace was cropped to 10-15 time points before the maximum peak in sensor readout, up to the maximum peak. The abrupt changes in sensor readout were identified using a change point algorithm^80, 81^ in Matlab called “findchangepts”, where abrupt changes in the slope were identified using the specifier, “linear’. A similar process was done to identify points of stoppage of cell elongation.

### Bistability Phase Plots

The mNG tag was used at the C-terminus of the Cdc13-Cdc2 fusion as it has better photophysical properties compared with sfGFP and allowed use of the YFP channel where there is less cellular autofluorescence^67, 77^. Experiments for Fig.4 and Extended Data Fig.4 were done using CellASIC ONIX V1 microfluidics platform using Y04C microfluidics plates (Merck), at 32° in YE4S, as they are the optimal conditions used for Cdc13-Cdc2AF^82^. All 4 experimental conditions (Cdc13-Cdc2-mNG + NucCDK/CytCDK V2-mScarletI, Cdc13-Cdc2AF-mNG + NucCDK/CytCDK V2-mScarletI) were done on the same plate using a flow rate of 2.5psi. 50uL of culture of an OD 0.2-0.3 was added to plate. Images were acquired every 5 minutes.

Phase plots were plotted after lightly smoothing the raw data using the matlab function, “smoothdata”, with the algorithm, “sgolay”, to reduce noise and improve the quality of the phase plots.

## Data and Code Availability

All scripts, strains, data, and plasmids, available upon request.

Analysis scripts can be found at: https://github.com/nkapadia27/Spatiotemporal-

Orchestration-of-Mitosis. DOI: 10.5281/zenodo.11072087.

## Acknowledgments

We thank T.Zeisner, J.Curran, and J.Greenwood for comments on the manuscript. This work was supported by the Francis Crick Institute, which receives core funding from Cancer Research UK (CC2003), the UK Medical Research Council (CC2003), and the Wellcome Trust (CC2003). For the purpose of Open Access, the author has applied a CC•BY copyright licence to any author accepted manuscript version arising from this submission. In addition, this work was supported by Wellcome Trust grants to P.N. (grant number 214183), and the Breast Cancer Research Foundation (BCRF-23-117). N.K was supported by an EMBO postdoctoral fellowship (EMBO ALTF 705-202) and a HFSP postdoctoral fellowship (HFSP LT000587/2021-L).

## Contributions

N.K. initiated the study. N.K designed and performed all the experiments. N.K. generated the strains and plasmids. N.K performed the image and data analysis including writing custom scripts. N.K. and P.N. wrote the manuscript.

---

**Extended Data Figure 1.**
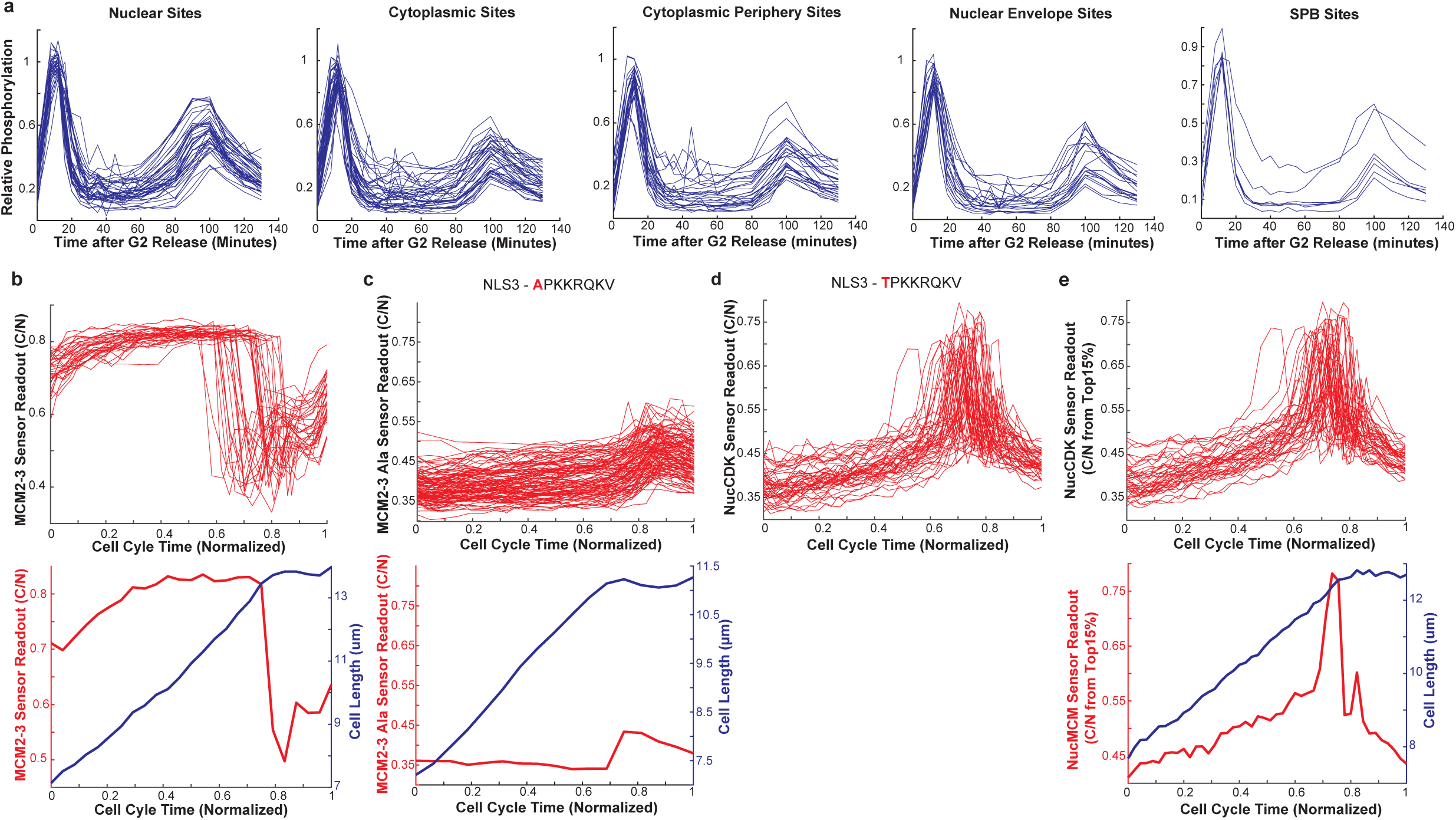
a) Profiles of individual late sites in the nucleus (*n* = 45 sites), cytoplasm (*n* = 41 sites), cytoplasmic periphery (*n* = 27 sites), nuclear envelope (*n* = 20 sites), and SPB (*n* = 7 sites). (b) Top: Traces of the original MCM2-3 sensor tagged with mScarletI, using the cytoplasmic to nuclear ratio as a measure of sensor readout, taken every 10 minutes, in EMM media. The mean top 15% of pixels were used as a measure for mean nuclear intensity, while the mean of bottom 85% of pixels was used as a measure for mean cytoplasmic intensity. *n* = 35 cells. Bottom: Representative time trace. c) Top: MCM2-3-Ala-mScarletI traces, taken every 5 minutes, in EMM media. Nuclear masks were obtained using MCM2-3-Ala-mScarletI. Y-axis scaled to MCM2-3 readout, shown in a). Bottom: Representative time trace of MCM2-3-Ala. d) NucCDK traces, where MCM2-3-Ala-mNG was used as a nuclear mask, taken every 5 minutes in EMM media. *n* = 42 cells. e) NucCDK time traces where the top 15% method was used with representative time trace (same cell as Figure 1c) shown on the bottom. *n* = 42 cells.

**Extended Data Figure 2.**
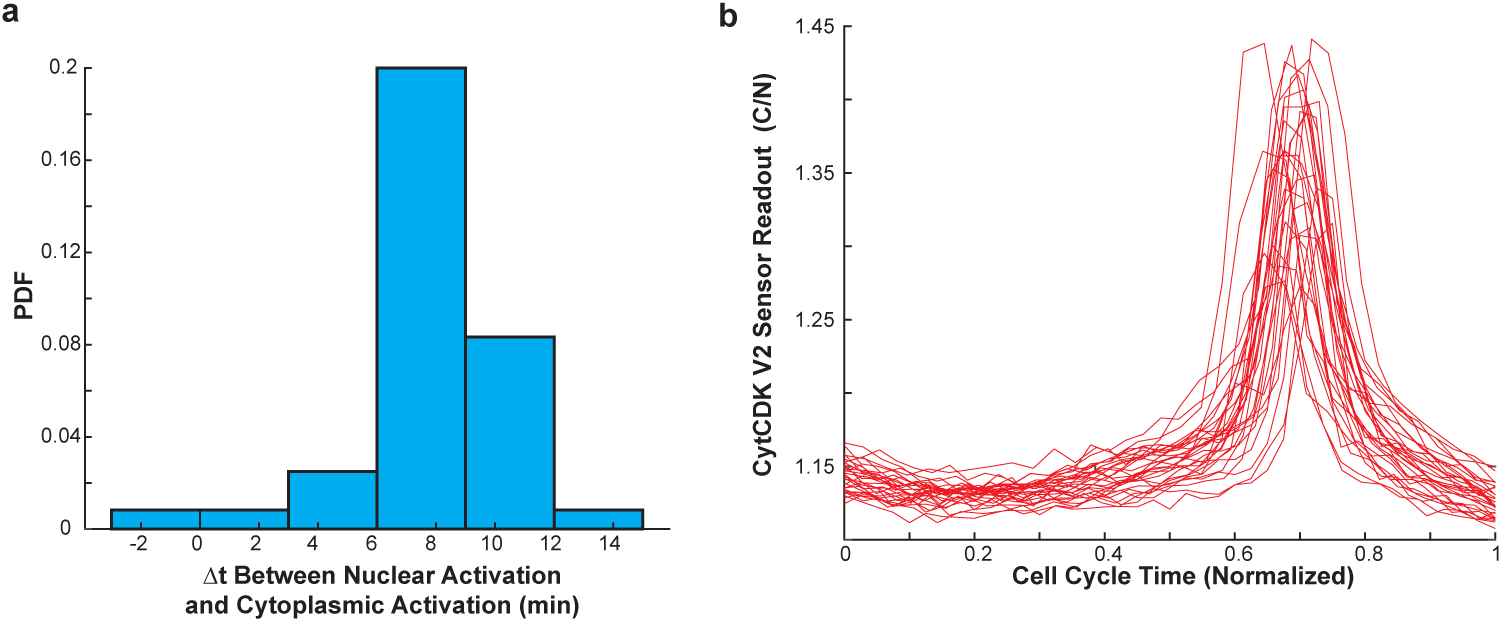
a) Histogram of the delay from nuclear activation to cytoplasmic activation of dual sensor strain with NucCDK-mScarletI and CytCDK-mNG using 3 minute intervals, in YE4S. Bin widths were set to time interval. *n* = 40 cells. b)Representative traces of CytCDK V2 from the dual sensor strain with Nuc-CDK-mScarletI. *n* = 30 cells.

**Extended Data Figure 3.**
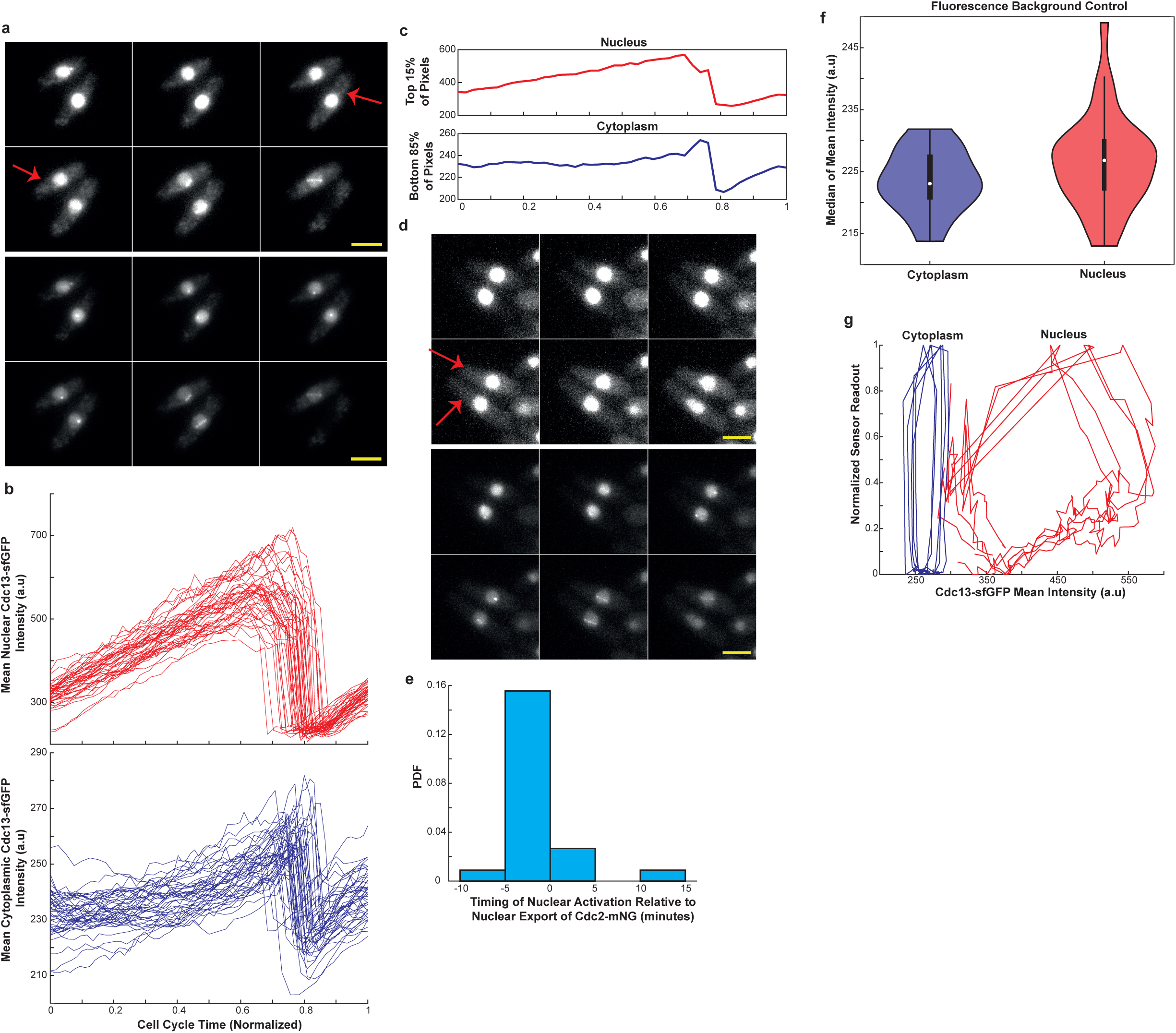
a) Top) Montage of Cdc13-sfGFP showing cells undergoing nuclear export. Images taken every 5 minutes, and each image represents 5 minutes. Scale bar = 5µm. Contrast was adjusted for better visualization of export. Red arrows mark cells where export was visually seen. Bottom) same as top but with contrast not enhanced. Cells were imaged using EMM media. b) Single cell traces showing mean intensities in the nucleus (top) and cytoplasm (bottom) of Cdc13-sfGFP. Data was collected every 5 minutes. MCM2-3-Ala-mScarletI was used as a nuclear mask. *n* = 45 cells. c) Representative trace of mean intensities of Cdc13-sfGFP in the nucleus and cytoplasm using top15% and bottom 85% of pixels, respectively. Same cell as in Figure 3b. d) Top) Montage of Cdc2-mNG taken every 5 minutes, showing two cells going through nuclear export. Each image is 5 minutes. Scale bar = 5µm. Contrast was adjusted for better visualization of export.Figure 3c but with contrast not enhanced. Bottom) Same as top but contrast not enhanced. Cells were imaged using EMM media. e) Histogram of nuclear activation using NucCDK-mScarletI, with repect to Cdc2-mNG nuclear export. Data was collected every 5 minutes. *n* = 45 cells. Bin width was set to time interval of acquisition (5 minutes). f) Background control of imaging in GFP channel with MCM2-3-Ala-mScarletI under same conditions used Fig 3h. Shown is the violin plot. The median value of the mean intensity of the nucleus and cytoplasm over the cell cycle was used. *n* = 33 cells. g) Phase plots of nucleus (mean nuclear Cdc13-sfGFP intensity and NucCDK readout) and cytoplasm (mean cytoplasmic Cdc13-sfGFP intensity and CytCDK readout), plotted on the same phase space. Sensor readouts were min max normalized for comparison. 5 cell traces are shown for each cellular compartment.

**Extended Data Figure 4.**
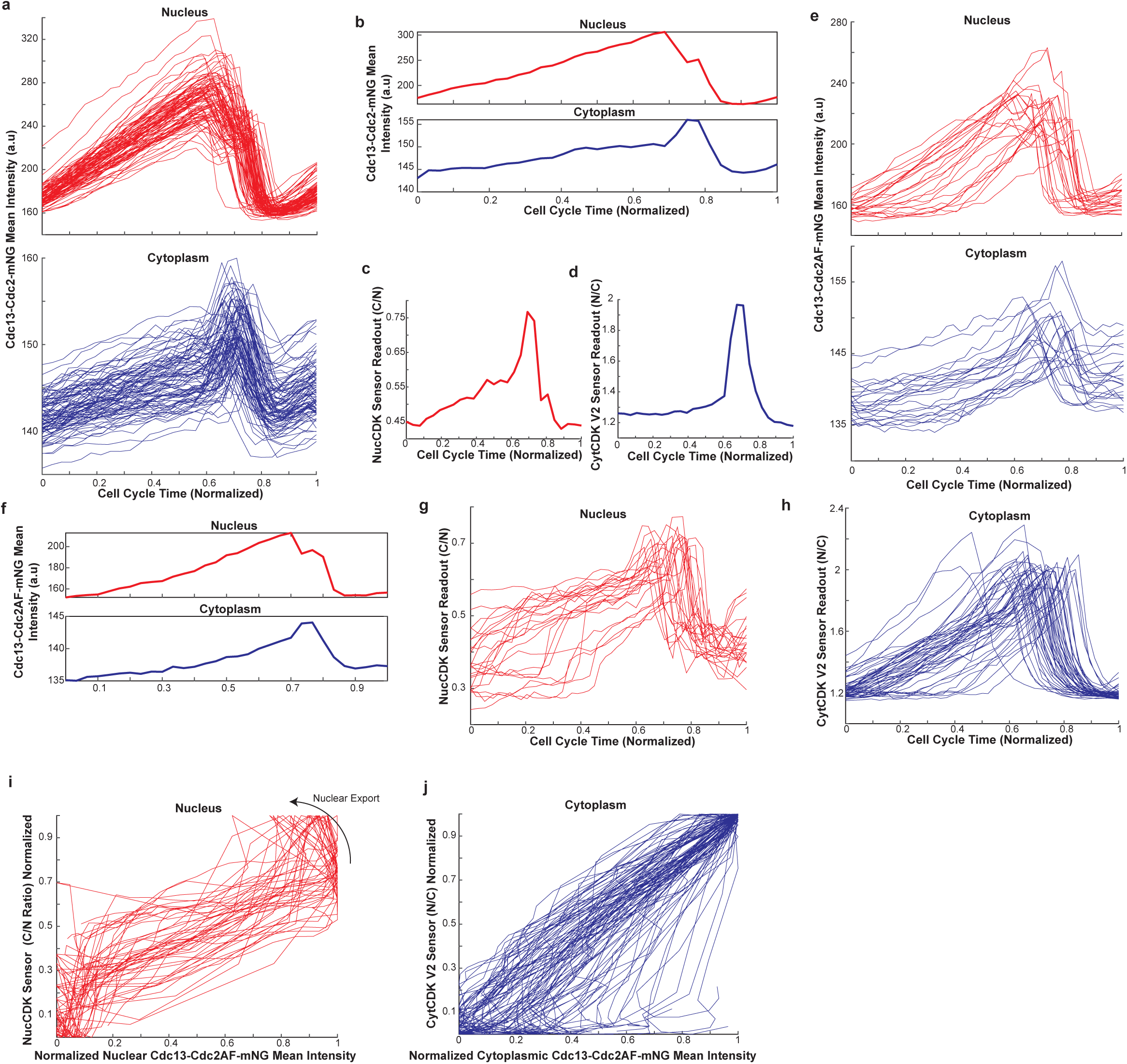
a) Single-cell traces of Cdc13-Cdc2-mNG mean intensities in nucleus (top) and cytoplasm (bottom). *n* = 89 cells. b) Representative trace of Cdc13-Cdc2-mNG mean intensities in the nucleus and cytoplasm. c) Representative trace of NucCDK-mScarletI in Cdc13-Cdc2-mNG background. d) Representative trace of CytCDK V2-mScarletI in Cdc13-Cdc2-mNG background. e) Single-cell traces of Cdc13-Cdc2AF-mNG mean intensities in nucleus (top) and cytoplasm (bottom). *n* = 24 cells. f) Representative trace of Cdc13-Cdc2AF-mNG mean intensities in the nucleus and cytoplasm. g) Single-cell traces of NucCDK sensor readout in Cdc13-Cdc2AF-mNG background. *n* = 24 cells. h) Single cell traces of CytCDK V2 sensor readout in Cdc13-Cdc2AF-mNG background. *n* = 49 cells. i) Min max normalized phase plots of the nucleus in the Cdc13-Cdc2AF-mNG background. *n* = 24 cells j) Min max normalized phase plots of the cytoplasm in the Cdc13-Cdc2AF-mNG background. *n* = 49 cells.

**Extended Data Figure 5.**
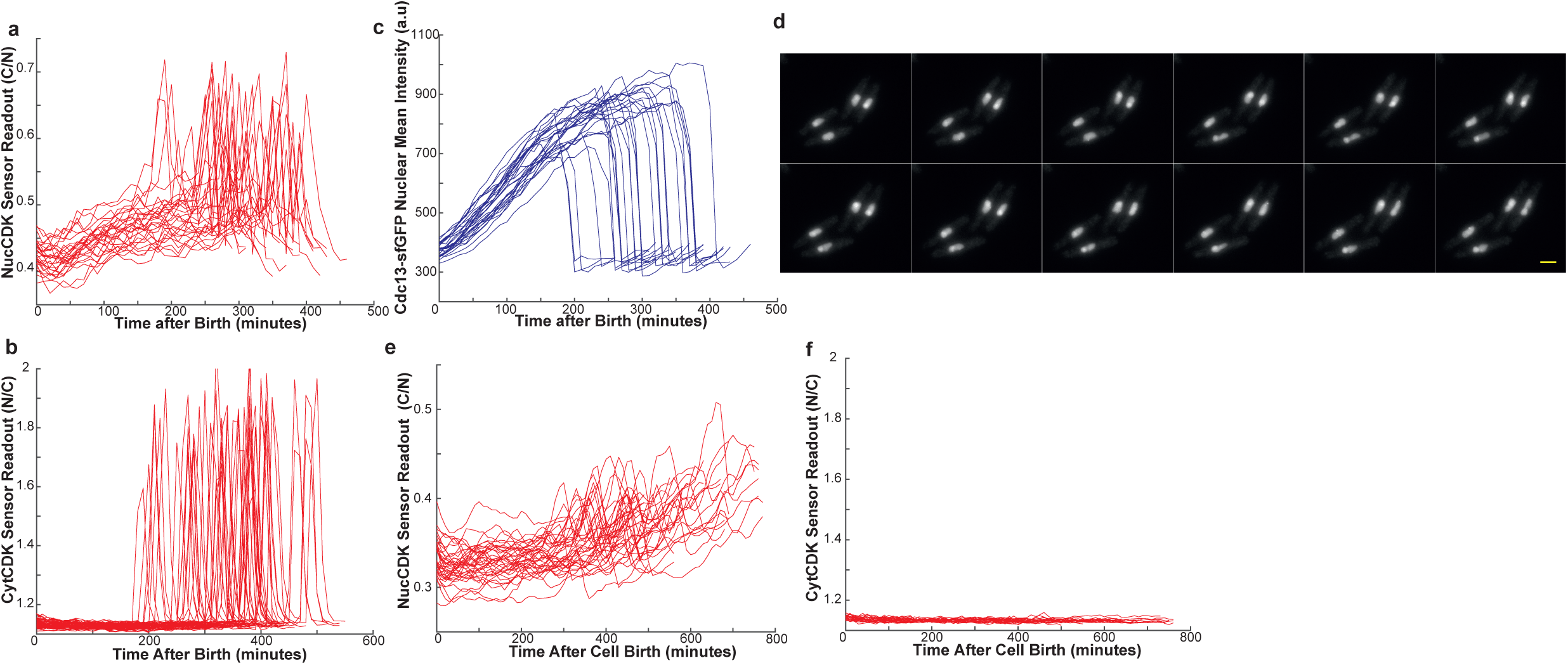
a) Single-cell traces of NucCDK readout in Cdc13^WT^-sfGFP background, in EMM + thiamine to repress the second copy of Cdc13^WT^. Images were acquired every 10 minutes. *n* = 29 cells. b) Single-cell traces of CytCDK readout in Cdc13^WT^-sfGFP in EMM + thiamine to repress the second copy of Cdc13^WT^. Images were acquired every 10 minutes. *n* = 58 cells. c) Single-cell traces of Cdc13-sfGFP mean nuclear intensity in Cdc13^WT^-sfGFP background, where the second copy of Cdc13^WT^ was repressed. *n* = 29 cells. d) Montage of Cdc13^HPM^-sfGFP. Images taken every 10 minutes, with each image representing 10 minutes. Scale bar = 5µm. e) Single-cell traces of NucCDK in Cdc13^HPM^-sfGFP background. *n* = 36 cells. f)Traces of CytCDK readout in Cdc13^HPM^-sfGFP background. Y-axis scaled based on minimum and maximum sensor readout in Cdc13^WT^ -sfGFP background shown in b). *n* = 23 cells.

## References

1 Grallert, A. et al. Centrosomal MPF triggers the mitotic and morphogenetic switches of fission yeast. Nat Cell Biol 15, 88–95, doi:10.1038/ncb2633 (2013).

2 Hachet, V., Canard, C. & Gonczy, P. Centrosomes promote timely mitotic entry in C. elegans embryos. Dev Cell 12, 531–541, doi:10.1016/j.devcel.2007.02.015 (2007).

3 Jackman, M., Lindon, C., Nigg, E. A. & Pines, J. Active cyclin B1-Cdk1 first appears on centrosomes in prophase. Nat Cell Biol 5, 143–148, doi:10.1038/ncb918 (2003).

4 Novak, B. & Tyson, J. J. Numerical analysis of a comprehensive model of M-phase control in Xenopus oocyte extracts and intact embryos. J Cell Sci 106 ( Pt 4), 1153–1168, doi:10.1242/jcs.106.4.1153 (1993).

5 Pomerening, J. R., Kim, S. Y. & Ferrell, J. E., Jr. Systems-level dissection of the cell-cycle oscillator: bypassing positive feedback produces damped oscillations. Cell 122, 565–578, doi:10.1016/j.cell.2005.06.016 (2005).

6 Pomerening, J. R., Sontag, E. D. & Ferrell, J. E., Jr. Building a cell cycle oscillator: hysteresis and bistability in the activation of Cdc2. Nat Cell Biol 5, 346–351, doi:10.1038/ncb954 (2003).

7 Rombouts, J. & Gelens, L. Dynamic bistable switches enhance robustness and accuracy of cell cycle transitions. PLoS Comput Biol 17, e1008231, doi:10.1371/journal.pcbi.1008231 (2021).

8 Sha, W. et al. Hysteresis drives cell-cycle transitions in Xenopus laevis egg extracts. Proc Natl Acad Sci U S A 100, 975–980, doi:10.1073/pnas.0235349100 (2003).

9 Dantas, M., Lima, J. T. & Ferreira, J. G. Nucleus-Cytoskeleton Crosstalk During Mitotic Entry. Front Cell Dev Biol 9, 649899, doi:10.3389/fcell.2021.649899 (2021).

10 Nurse, P. Universal control mechanism regulating onset of M-phase. Nature 344, 503–508, doi:10.1038/344503a0 (1990).

11 Swaffer, M. P., Jones, A. W., Flynn, H. R., Snijders, A. P. & Nurse, P. CDK Substrate Phosphorylation and Ordering the Cell Cycle. Cell 167, 1750–1761 e1716, doi:10.1016/j.cell.2016.11.034 (2016).

12 Hegarat, N. et al. Cyclin A triggers Mitosis either via the Greatwall kinase pathway or Cyclin B. EMBO J 39, e104419, doi:10.15252/embj.2020104419 (2020).

13 Gong, D. et al. Cyclin A2 regulates nuclear-envelope breakdown and the nuclear accumulation of cyclin B1. Curr Biol 17, 85–91, doi:10.1016/j.cub.2006.11.066 (2007).

14 Lindqvist, A., Rodriguez-Bravo, V. & Medema, R. H. The decision to enter mitosis: feedback and redundancy in the mitotic entry network. J Cell Biol 185, 193–202, doi:10.1083/jcb.200812045 (2009).

15 Pagliuca, F. W. et al. Quantitative proteomics reveals the basis for the biochemical specificity of the cell-cycle machinery. Mol Cell 43, 406–417, doi:10.1016/j.molcel.2011.05.031 (2011).

16 Deibler, R. W. & Kirschner, M. W. Quantitative reconstitution of mitotic CDK1 activation in somatic cell extracts. Mol Cell 37, 753–767, doi:10.1016/j.molcel.2010.02.023 (2010).

17 Gong, D. & Ferrell, J. E., Jr. The roles of cyclin A2, B1, and B2 in early and late mitotic events. Mol Biol Cell 21, 3149–3161, doi:10.1091/mbc.E10-05-0393 (2010).

18 Gavet, O. & Pines, J. Activation of cyclin B1-Cdk1 synchronizes events in the nucleus and the cytoplasm at mitosis. J Cell Biol 189, 247–259, doi:10.1083/jcb.200909144 (2010).

19 Gavet, O. & Pines, J. Progressive activation of CyclinB1-Cdk1 coordinates entry to mitosis. Dev Cell 18, 533–543, doi:10.1016/j.devcel.2010.02.013 (2010).

20 Nigg, E. A. Cellular substrates of p34(cdc2) and its companion cyclin-dependent kinases. Trends Cell Biol 3, 296–301, doi:10.1016/0962-8924(93)90011-o (1993).

21 Welburn, J. P. et al. How tyrosine 15 phosphorylation inhibits the activity of cyclin-dependent kinase 2-cyclin A. J Biol Chem 282, 3173–3181, doi:10.1074/jbc.M609151200 (2007).

22 Murray, A. W. & Kirschner, M. W. Cyclin synthesis drives the early embryonic cell cycle. Nature 339, 275–280, doi:10.1038/339275a0 (1989).

23 Strausfeld, U. et al. Dephosphorylation and activation of a p34cdc2/cyclin B complex in vitro by human CDC25 protein. Nature 351, 242–245, doi:10.1038/351242a0 (1991).

24 Russell, P. & Nurse, P. Negative regulation of mitosis by wee1+, a gene encoding a protein kinase homolog. Cell 49, 559–567, doi:10.1016/0092-8674(87)90458-2 (1987).

25 Millar, J. B., McGowan, C. H., Lenaers, G., Jones, R. & Russell, P. p80cdc25 mitotic inducer is the tyrosine phosphatase that activates p34cdc2 kinase in fission yeast. EMBO J 10, 4301–4309, doi:10.1002/j.1460-2075.1991.tb05008.x (1991).

26 Gautier, J., Solomon, M. J., Booher, R. N., Bazan, J. F. & Kirschner, M. W. cdc25 is a specific tyrosine phosphatase that directly activates p34cdc2. Cell 67, 197–211, doi:10.1016/0092-8674(91)90583-k (1991).

27 Parker, L. L. & Piwnica-Worms, H. Inactivation of the p34cdc2-cyclin B complex by the human WEE1 tyrosine kinase. Science 257, 1955–1957, doi:10.1126/science.1384126 (1992).

28 Kim, S. Y. & Ferrell, J. E., Jr. Substrate competition as a source of ultrasensitivity in the inactivation of Wee1. Cell 128, 1133–1145, doi:10.1016/j.cell.2007.01.039 (2007).

29 Trunnell, N. B., Poon, A. C., Kim, S. Y. & Ferrell, J. E., Jr. Ultrasensitivity in the Regulation of Cdc25C by Cdk1. Mol Cell 41, 263–274, doi:10.1016/j.molcel.2011.01.012 (2011).

30 Hoffmann, I., Clarke, P. R., Marcote, M. J., Karsenti, E. & Draetta, G. Phosphorylation and activation of human cdc25-C by cdc2--cyclin B and its involvement in the self-amplification of MPF at mitosis. EMBO J 12, 53–63, doi:10.1002/j.1460-2075.1993.tb05631.x (1993).

31 McGowan, C. H. & Russell, P. Cell cycle regulation of human WEE1. EMBO J 14, 2166–2175, doi:10.1002/j.1460-2075.1995.tb07210.x (1995).

32 Pomerening, J. R., Ubersax, J. A. & Ferrell, J. E., Jr. Rapid cycling and precocious termination of G1 phase in cells expressing CDK1AF. Mol Biol Cell 19, 3426–3441, doi:10.1091/mbc.e08-02-0172 (2008).

33 Pines, J. & Hunter, T. Human cyclins A and B1 are differentially located in the cell and undergo cell cycle-dependent nuclear transport. J Cell Biol 115, 1–17, doi:10.1083/jcb.115.1.1 (1991).

34 Santos, S. D., Wollman, R., Meyer, T. & Ferrell, J. E., Jr. Spatial positive feedback at the onset of mitosis. Cell 149, 1500–1513, doi:10.1016/j.cell.2012.05.028 (2012).

35 De Souza, C. P., Ellem, K. A. & Gabrielli, B. G. Centrosomal and cytoplasmic Cdc2/cyclin B1 activation precedes nuclear mitotic events. Exp Cell Res 257, 11–21, doi:10.1006/excr.2000.4872 (2000).

36 Arquint, C., Gabryjonczyk, A. M. & Nigg, E. A. Centrosomes as signalling centres. Philos Trans R Soc Lond B Biol Sci 369, doi:10.1098/rstb.2013.0464 (2014).

37 Nolet, F. E. et al. Nuclei determine the spatial origin of mitotic waves. Elife 9, doi:10.7554/eLife.52868 (2020).

38 Afanzar, O., Buss, G. K., Stearns, T. & Ferrell, J. E., Jr. The nucleus serves as the pacemaker for the cell cycle. Elife 9, doi:10.7554/eLife.59989 (2020).

39 Heald, R., McLoughlin, M. & McKeon, F. Human wee1 maintains mitotic timing by protecting the nucleus from cytoplasmically activated Cdc2 kinase. Cell 74, 463–474, doi:10.1016/0092-8674(93)80048-j (1993).

40 Koivomagi, M. et al. Dynamics of Cdk1 substrate specificity during the cell cycle. Mol Cell 42, 610–623, doi:10.1016/j.molcel.2011.05.016 (2011).

41 Ord, M. et al. Multisite phosphorylation code of CDK. Nat Struct Mol Biol 26, 649–658, doi:10.1038/s41594-019-0256-4 (2019).

42 Ord, M. et al. Proline-Rich Motifs Control G2-CDK Target Phosphorylation and Priming an Anchoring Protein for Polo Kinase Localization. Cell Rep 31, 107757, doi:10.1016/j.celrep.2020.107757 (2020).

43 Coudreuse, D. & Nurse, P. Driving the cell cycle with a minimal CDK control network. Nature 468, 1074–1079, doi:10.1038/nature09543 (2010).

44 Booher, R. N., Alfa, C. E., Hyams, J. S. & Beach, D. H. The fission yeast cdc2/cdc13/suc1 protein kinase: regulation of catalytic activity and nuclear localization. Cell 58, 485–497, doi:10.1016/0092-8674(89)90429-7 (1989).

45 Lee, M. G. & Nurse, P. Complementation used to clone a human homologue of the fission yeast cell cycle control gene cdc2. Nature 327, 31–35, doi:10.1038/327031a0 (1987).

46 Nurse, P. Genetic control of cell size at cell division in yeast. Nature 256, 547–551, doi:10.1038/256547a0 (1975).

47 Basu, S. et al. The Hydrophobic Patch Directs Cyclin B to Centrosomes to Promote Global CDK Phosphorylation at Mitosis. Curr Biol 30, 883–892 e884, doi:10.1016/j.cub.2019.12.053 (2020).

48 Piel, M. & Tran, P. T. Cell shape and cell division in fission yeast. Curr Biol 19, R823–827, doi:10.1016/j.cub.2009.08.012 (2009).

49 Nurse, P., Thuriaux, P. & Nasmyth, K. Genetic control of the cell division cycle in the fission yeast Schizosaccharomyces pombe. Mol Gen Genet 146, 167–178, doi:10.1007/BF00268085 (1976).

50 Patterson, J. O., Basu, S., Rees, P. & Nurse, P. CDK control pathways integrate cell size and ploidy information to control cell division. Elife 10, doi:10.7554/eLife.64592 (2021).

51 Curran, S., Dey, G., Rees, P. & Nurse, P. A quantitative and spatial analysis of cell cycle regulators during the fission yeast cycle. Proc Natl Acad Sci U S A 119, e2206172119, doi:10.1073/pnas.2206172119 (2022).

52 Grallert, A. et al. Removal of centrosomal PP1 by NIMA kinase unlocks the MPF feedback loop to promote mitotic commitment in S. pombe. Curr Biol 23, 213–222, doi:10.1016/j.cub.2012.12.039 (2013).

53 MacIver, F. H., Tanaka, K., Robertson, A. M. & Hagan, I. M. Physical and functional interactions between polo kinase and the spindle pole component Cut12 regulate mitotic commitment in S. pombe. Genes Dev 17, 1507–1523, doi:10.1101/gad.256003 (2003).

54 Mulvihill, D. P., Petersen, J., Ohkura, H., Glover, D. M. & Hagan, I. M. Plo1 kinase recruitment to the spindle pole body and its role in cell division in Schizosaccharomyces pombe. Mol Biol Cell 10, 2771–2785, doi:10.1091/mbc.10.8.2771 (1999).

55 Tanaka, K. et al. The role of Plo1 kinase in mitotic commitment and septation in Schizosaccharomyces pombe. EMBO J 20, 1259–1270, doi:10.1093/emboj/20.6.1259 (2001).

56 Roberts, E. L. et al. CDK activity at the centrosome regulates the cell cycle. Cell Rep 43, 114066, doi:10.1016/j.celrep.2024.114066 (2024).

57 Ferrell, J. E. & Xiong, W. Bistability in cell signaling: How to make continuous processes discontinuous, and reversible processes irreversible. Chaos 11, 227–236, doi:10.1063/1.1349894 (2001).

58 Yao, G., Lee, T. J., Mori, S., Nevins, J. R. & You, L. A bistable Rb-E2F switch underlies the restriction point. Nat Cell Biol 10, 476–482, doi:10.1038/ncb1711 (2008).

59 Tuck, C., Zhang, T., Potapova, T., Malumbres, M. & Novak, B. Robust mitotic entry is ensured by a latching switch. Biol Open 2, 924–931, doi:10.1242/bio.20135199 (2013).

60 Ferrell, J. E., Jr. How regulated protein translocation can produce switch-like responses. Trends Biochem Sci 23, 461–465, doi:10.1016/s0968-0004(98)01316-4 (1998).

61 Castedo, M. et al. Cell death by mitotic catastrophe: a molecular definition. Oncogene 23, 2825–2837, doi:10.1038/sj.onc.1207528 (2004).

62 Lemmens, B. et al. DNA Replication Determines Timing of Mitosis by Restricting CDK1 and PLK1 Activation. Mol Cell 71, 117–128 e113, doi:10.1016/j.molcel.2018.05.026 (2018).

63 Matsuyama, M. et al. Nuclear Chk1 prevents premature mitotic entry. J Cell Sci 124, 2113–2119, doi:10.1242/jcs.086488 (2011).

64 Wood, E. & Nurse, P. Sizing up to divide: mitotic cell-size control in fission yeast. Annu Rev Cell Dev Biol 31, 11–29, doi:10.1146/annurev-cellbio-100814-125601 (2015).

65 Tsai, T. Y. et al. Robust, tunable biological oscillations from interlinked positive and negative feedback loops. Science 321, 126–129, doi:10.1126/science.1156951 (2008).

## Methods References

66 Matsuyama, A., Shirai, A. & Yoshida, M. A series of promoters for constitutive expression of heterologous genes in fission yeast. Yeast 25, 371–376, doi:10.1002/yea.1593 (2008).

67 Shaner, N. C. et al. A bright monomeric green fluorescent protein derived from Branchiostoma lanceolatum. Nat Methods 10, 407–409, doi:10.1038/nmeth.2413 (2013).

68 Bindels, D. S. et al. mScarlet: a bright monomeric red fluorescent protein for cellular imaging. Nat Methods 14, 53–56, doi:10.1038/nmeth.4074 (2017).

69 Liku, M. E., Nguyen, V. Q., Rosales, A. W., Irie, K. & Li, J. J. CDK phosphorylation of a novel NLS-NES module distributed between two subunits of the Mcm2-7 complex prevents chromosomal rereplication. Mol Biol Cell 16, 5026–5039, doi:10.1091/mbc.e05-05-0412 (2005).

70 Godfrey, M. et al. PP2A(Cdc55) Phosphatase Imposes Ordered Cell-Cycle Phosphorylation by Opposing Threonine Phosphorylation. Mol Cell 65, 393–402 e393, doi:10.1016/j.molcel.2016.12.018 (2017).

71 Sugiyama, H., Goto, Y., Kondo, Y., Coudreuse, D. & Aoki, K. Live-cell imaging defines a threshold in CDK activity at the G2/M transition. Dev Cell 59, 545–557 e544, doi:10.1016/j.devcel.2023.12.014 (2024).

72 Moreno, S., Klar, A. & Nurse, P. Molecular genetic analysis of fission yeast Schizosaccharomyces pombe. Methods Enzymol 194, 795–823, doi:10.1016/0076-6879(91)94059-l (1991).

73 Kamenz, J., Mihaljev, T., Kubis, A., Legewie, S. & Hauf, S. Robust Ordering of Anaphase Events by Adaptive Thresholds and Competing Degradation Pathways. Mol Cell 60, 446–459, doi:10.1016/j.molcel.2015.09.022 (2015).

74 Kapadia, N. et al. Processive Activity of Replicative DNA Polymerases in the Replisome of Live Eukaryotic Cells. Mol Cell 80, 114–126 e118, doi:10.1016/j.molcel.2020.08.014 (2020).

75 Laplante, C., Huang, F., Bewersdorf, J. & Pollard, T. D. High-Speed Super-Resolution Imaging of Live Fission Yeast Cells. Methods Mol Biol 1369, 45–57, doi:10.1007/978-1-4939-3145-3_4 (2016).

76 Edelstein, A. D. et al. Advanced methods of microscope control using muManager software. J Biol Methods 1, doi:10.14440/jbm.2014.36 (2014).

77 Davidson, R., Liu, Y., Gerien, K. S. & Wu, J. Q. Real-Time Visualization and Quantification of Contractile Ring Proteins in Single Living Cells. Methods Mol Biol 1369, 9–23, doi:10.1007/978-1-4939-3145-3_2 (2016).

78 Schindelin, J. et al. Fiji: an open-source platform for biological-image analysis. Nat Methods 9, 676–682, doi:10.1038/nmeth.2019 (2012).

79 Chalfoun, J. et al. Lineage mapper: A versatile cell and particle tracker. Sci Rep 6, 36984, doi:10.1038/srep36984 (2016).

80 Killick, R., Fearnhead, P. & Eckley, I. A. Optimal Detection of Changepoints With a Linear Computational Cost. Journal of the American Statistical Association 107, 1590–1598, doi:10.1080/01621459.2012.737745 (2012).

81 Lavielle, M. Using penalized contrasts for the change-point problem. Signal Processing 85, 1501–1510, 10.1016/j.sigpro.2005.01.012 (2005).

82 Scotchman, E., Kume, K., Navarro, F. J. & Nurse, P. Identification of mutants with increased variation in cell size at onset of mitosis in fission yeast. J Cell Sci 134, doi:10.1242/jcs.251769 (2021).

